# Founder cell configuration drives competitive outcome within colony biofilms

**DOI:** 10.1101/2021.07.08.451560

**Authors:** Lukas Eigentler, Margarita Kalamara, Graeme Ball, Cait E. MacPhee, Nicola R. Stanley-Wall, Fordyce A. Davidson

**Affiliations:** Division of Molecular Microbiology, School of Life Sciences, University of Dundee, Dundee DD1 5EH, United Kingdom; Division of Mathematics, School of Science and Engineering, University of Dundee, Dundee DD1 4HN, United Kingdom; Dundee Imaging Facility, School of Life Sciences, University of Dundee, Dundee DD1 5HN, United Kingdom; School of Physics and Astronomy, The University of Edinburgh, Edinburgh EH9 3FD, UK

**Author notes:** Joint Corresponding Authors.

## Abstract

Bacteria typically form dense communities called biofilms, where cells are embedded in a self-produced extracellular matrix. Competitive interactions between strains within the biofilm context are studied due to their potential applications in biological, medical, and industrial systems. Combining mathematical modelling with experimental assays, we reveal that the spatial structure and the competitive dynamics within biofilms are significantly affected by the location and density of founder cells. Using an isogenic pair of Bacillus subtilis strains, we show that the observed spatial structure and relative strain biomass in a mature biofilm can be mapped directly to the locations of founder cells. Moreover, we define a predictor of competitive outcome that accurately forecasts relative abundance of strains based solely on the founder cells’ access to free space. Consequently, we reveal that variability of competitive outcome in biofilms inoculated at low founder density is a natural consequence of the random positioning of founding cells in the inoculum. Extending our study to non-isogenic strain pairs of B. subtilis, we show that even for strains with different antagonistic strengths, a race for space remains the dominant mode of competition in biofilms inoculated at low founder densities. Our results highlight the importance of spatial dynamics on competitive interactions within biofilms and hence to related applications.

## 1. Introduction

Biofilms are consortia of microorganisms (1): cells embedded in a self-produced extracellular matrix typically comprising extracellular polysaccharides, proteins, DNA, and components of lysed cells (1–5). Many natural, industrial, and medical environments are significantly affected by biofilm formation. For example, biofilms are used in wastewater treatment (6), are fundamental to the functioning of microbiota in the human gastrointestinal tract (7), and are central to global biogeochemical cycling (8, 9). However, from a human perspective, the role of biofilms is not exclusively positive, as they are a known cause of the majority of chronic infections (10) and medical device fouling (11).

Many *in vitro* models have been developed to study diverse biofilms and the complexity of the model set-up, and the ability of each system to mimic physiologically relevant conditions is highly varied (3). One simple, but widely used model is the colony biofilm model. In this system, founding cells are deposited on an agar-solidified growth medium and the architecturally complex macroscale structure that develops is thus easily examined (3). The colony biofilm model has proven effective in revealing regulatory pathways involved in controlling biofilm formation and the production of molecules found in the biofilm matrix. Hence, it has been implemented widely across many different microorganisms (1, 3, 12).

Increasingly, the colony biofilm model is being exploited to explore competitive dynamics within single and mixed species biofilms (13–19). Many competitive mechanisms fundamentally depend on spatial co-location of strains, with spatial segregation within multi-strain biofilms offering protection.(14, 17, 20, 21). Hence, a necessary precursor to a full understanding of the effects of interbacterial killing mechanisms on competitive dynamics within biofilms is a comprehensive understanding of the impact of spatial dynamics. One way to artificially impose spatial segregation in biofilms is the deposition of founder cells in a prescribed spatial structure (20, 22–25). However, spatial segregation within multi-strain biofilms can also arise from well-mixed founder cells, particularly if the density of founder cells is low (14, 16, 17, 21, 26). The observed increase of spatial segregation with decreasing founder density is typically attributed to increased separation of founder cells. The hypothesis is that this initial separation allows for the establishment of distinct, single-strain spatial patches before these expanding micro-colonies encounter each other (14, 17). Despite a growing body of work in this area, the mechanisms by which competition for space is affected by the location and density of founder cells are still unclear. In particular, why small numbers of founder cells lead to spatial segregation in mature biofilms, and impact competitive dynamics underpinned by killing mechanisms, remain open questions.

Here, we elucidate the fundamental role of spatial dynamics in single-and dual-strain colony biofilms across a wide range of founder densities by combining mathematical modelling with an experimental co-culture colony biofilm assay using *Bacillus subtilis*, a well-understood Gram-positive bacterium. We first focussed on single-strain biofilms to restrict competitive dynamics between founder cells to those related only to space. We mapped parts of the mature biofilm to founder cells and showed that the random placement of founder cells significantly affected the biofilm’s spatial structure. This analysis led to the development of a predictor for competitive outcome whose accuracy was retained in an extension of our iterative approach to dual-strain biofilms. Implementing the predictor revealed that competition for space was the dominating mode of competition, even for strain pairs in which the strengths of antagonistic interactions were disproportionally skewed towards one strain.

## 2. Results

### A theoretical framework of interacting bacterial strains

We investigated the impact of founder cell locations on the structure of single-strain biofilms, using an iterative approach of mathematical modelling and experimental assays. The mathematical model was motivated by experimental assays used to establish colony biofilms on agar surfaces where the founding inoculum is placed on the surface of nutrient agar. Within the initial footprint, individual (or small clusters of) bacteria settle at random locations and grow over time into a mature structured macroscale community (Figure 1A). In the mathematical model all the founding cells have identical properties. However, to track the dynamics of biofilm growth we divided the founding cells into two groups, denoted by *B*_1_ (visualised in magenta colour) and *B*_2_ (visualised in green colour) (Figure 1B). Note that we refer to *B*_1_ and *B*_2_ as *strains* for brevity, even though they simulate two isogenic cell lineages which express different fluorescent proteins in a single-strain biofilm (Figure 1A). Growth of the strains was described by equations based on the spatially extended competitive Lotka-Volterra model. Suitably nondimensionalised (see Section S3), the model is given by

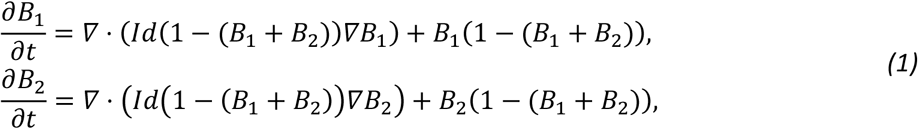

where, the variables 0 ≤ *B*_1_(*x*, *t*), *B*_2_(*x*, *t*) ≤ 1 denote the scaled densities of each strain, respectively at time *t* >0 (one nondimensional time unit corresponding to approx. 2.9 h) and at spatial position *x* ∈ Ω (one nondimensional space unit corresponding to approx. 0.15 mm). The spatial domain 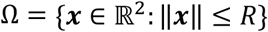 is a disk, representing the biofilm growth medium (Figure 1C).

**Figure 1:**
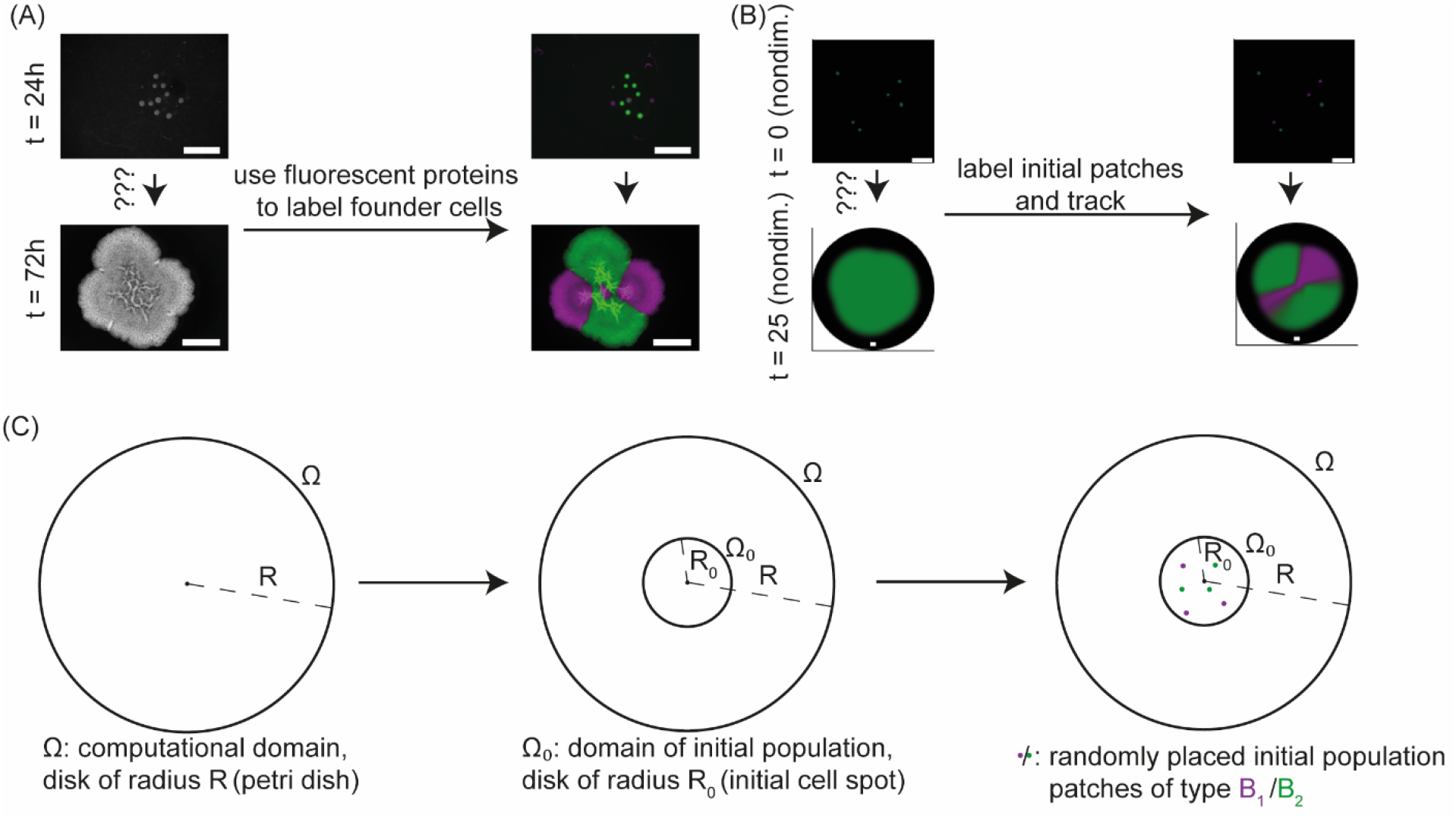
Experimental and modelling set-up. (A) An example of the experimental assay. Founder cells carry either a constitutively expressed copy of gfp (green) or mTagBFP (magenta). The bacteria were mixed in a 1:1 ratio and images taken after 24h and 72h of incubation. The initial founder density was approx. 10 CFUs. The scalebars are 5 mm long. (B) An example realisation of the mathematical model. In the right-hand plots green and magenta are used to differentiate two subsets of the initial patches (*t* = 0, top) and their subsequent development (*t* = 25, bottom). Black areas indicate the computational domain, *Ω*. The plot of initial condition is a blow-up of the centre of the whole domain. The scale bars represent 7 nondimensional space units. (C) Schematic of model initial condition. Initial populations (filled coloured circles) are placed in *Ω*_0_, a small subdomain of the whole computational domain *Ω* (both centred at the origin *O*).

The initial conditions of the theoretical framework were motivated by the random positions at which bacteria settle on the agar within the inoculum footprint (Figure 1A). In our theoretical framework, we represented the experimental inoculum footprint by the subdomain Ω_0_ = {*x* ∈ Ω: ||*x*|| < *R*_0_} (Figure 1C). We mimicked the random deposition of bacteria (see below) by randomly placing “microcolonies” within Ω_0_ at nodes of a triangulated spatial mesh of linear geometric order, used in the application of a finite element method to numerically solve the model equations (Figure 1B and 1C). Each initial microcolony was assumed to only contain one strain and to be at carrying capacity (i.e. *B*_1_ = 1 or *B*_2_ = 1 within each microcolony). Unless otherwise stated, we used an even number (*N*) of initial microcolonies and assigned exactly *N*/2 to each strain at random. At spatial locations other than the assigned microcolonies, both densities were set to zero.

The size of a spatial mesh element used in the model (approx. 0.008 *mm*^2^ in experimental parameters) was much larger than that of a single bacterial cell. This means that the initial conditions represented the experimental assays shortly after inoculation (typically after 24h of incubation), at which time each bacterium (or small cluster of bacteria) had formed a *distinct*, *spatially separated microcolony*. Hence, the number of *model microcolonies N* represents the number of bacteria used in the initial inoculum. Resolving the initial data at this spatial scale allowed analysis for 0 ≤ *N* ≤ 824. Using a selected set of numbers from that range was sufficient to capture clear trends (see below). The range covers biologically relevant founder densities, which generate mature colony biofilms with broadly similar morphologies (Figure S1). Additionally, to verify whether the observed trends could be extrapolated to *N* > 824, we represented high founder densities by piecewise spatially homogeneous initial conditions *B*_1_ = *B*_2_ = 0.5 in Ω_0_ and *B*_1_ = *B*_2_ = 0 otherwise.

The strains are assumed to grow logistically, with growth being limited by the total population, which cannot exceed unity (after nondimensionalisation). Moreover, spatial propagation is described by diffusion as is common (27). However, in our model, we employ a diffusion coefficient that decreases with increasing population size. This density dependence prevents merging of initially separated founding patches in the model and is invoked to capture experimental observations that indicate such colonies abut rather than merge on meeting (28, 29). The indicator function *Id* = 1 if *B*_1_ + *B*_2_ ≤ 1 and Id = 0 otherwise guarantees nonnegativity of the diffusion coefficients; this constrains the model to the physically relevant case and moreover ensures numerical stability during simulation.

Finally, we defined the *competitive outcome* score of the interaction to be the relative mass of strain *B*_1_ i.e. 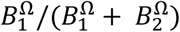 at the chosen end point (*t* = *T*) of our model simulation, where

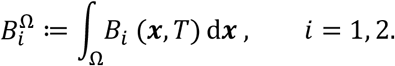

The competitive outcome score lies in the interval [0,1] with the value 0.5 signifying a 1:1 ratio between the strains. Note that we could swap the indices without loss of generality to equivalently define the competitive outcome to be the relative mass of strain *B*_2_ at the chosen end point.

### Low founder densities yield large variability in competitive outcomes

In the absence of spatial dynamics, the mathematical model predicts that the ratio between both strains would always remain constant 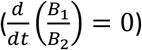 and therefore that the competitive outcome would completely be determined by the initial ratio. To test whether such a relationship continues to hold in the full, spatially extended system, we examined data from simulations over a test range of initial founding cell densities. The initial strain ratio was selected to be 1:1 for each test.

Model simulations using homogeneous initial conditions (representing high founder densities) consistently resulted in a *competitive outcome* score of 0.5 (i.e. strains in 1:1 ratio) and with the strains remaining homogeneously distributed in space across the colony (Figure 2A, Movie S1). By contrast, independent model realisations using a specified number of microcolonies placed at randomly chosen locations representing low (*N* = 6) and intermediate (*N* = 824) founder densities, revealed significant variation in competitive outcome (Figure 2B and 2C, Movies S2 and S3). To explore this observed variability in more detail, we employed a Monte Carlo approach. For each fixed number *N* within the selected set, 1000 independent model realisations were conducted. Data from these simulations revealed the competitive outcome score for each founder density to be normally distributed with mean 0.5. Moreover, the standard deviation decreased with increasing *N* (Figure 2D). (The exception is the case *N* = 2; see supplementary information for a discussion of this special case). Finally, our model predicts significant changes in the spatial organisation of the two strains within the biofilm in response to changing founder density (see also (14)). For high founder densities, isogenic strains are predicted to coexist homogenously (Figure 2A). However, as the founder density is decreased (decreasing *N*), the model predicts that homogeneous coexistence is gradually replaced by the formation of spatial sectors dominated by one strain or the other with full spatial segregation resulting from low founder densities (Figure 2B and 2C). This model prediction is consistent with previous studies, which highlight that the deposition of founder cells at low density can lead to spatial segregation in both single- and multi-strain biofilms (14, 16, 17, 21, 26).

**Figure 2:**
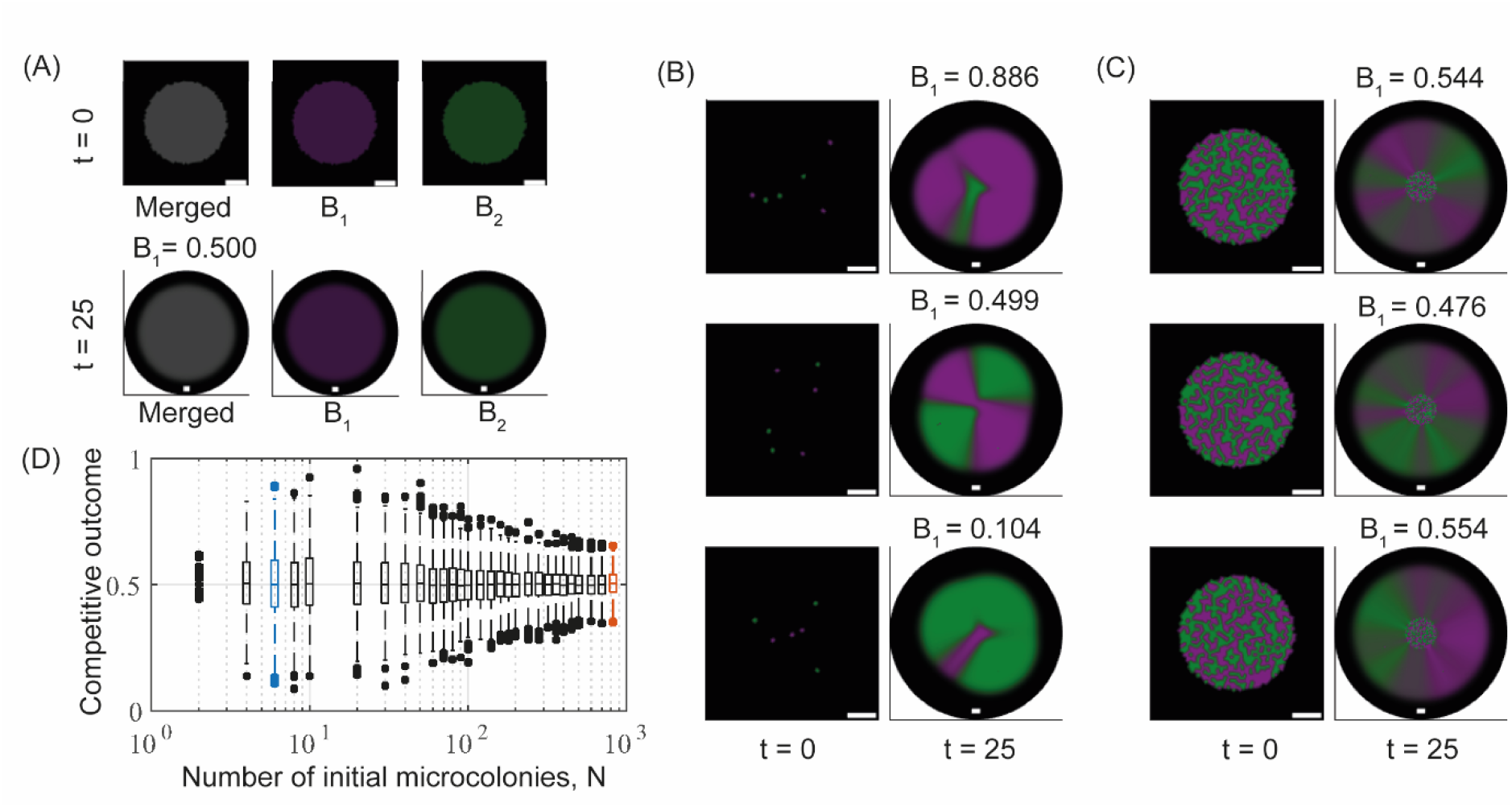
Variability in spatial structure at low founder density. (A-C) Example model realisations for different founder densities. All plots show the system’s initial conditions (*t* = 0) and the outcomes after 25 time units. Plots visualising the systems’ states at *t* = 0 show a blow-up of the subdomain *Ω*_0_; plots visualising outcomes at *t* = 25 show the full computational domain *Ω* (black background). The scalebars are seven unit lengths long. (A) the outcome of simulations initialised with piecewise spatially homogeneous populations representing high founder density. The “Merged” image channel shows both strains (grey colour corresponds to overlap); the *B*_1_(green) and *B*_2_ (magenta) channels only show single strain filters of the plot. (B) the range of outcomes observed for low founder density (number of initial cell patches *N* = 6). (C) more consistent outcomes for intermediate founder densities (*N* = 824). In (B, C), only the “Merged” channel is shown. (D) Variability in competitive outcome increases with decreasing founder density. Each boxplot contains data from 1000 model realisations. Blue and red boxplots correspond to the founder densities in (B) and (C), respectively.

### Access to free space determines competitive outcome

Next, we attempted to uncover the mechanism(s) by which low founder densities drive variability in competitive outcome. Motivated by (14), we first considered whether the initial separation between initial microcolonies of different types was the simple determinant. We did not find this to be the case for isogenic strain pairings (Figure S2). As an alternative, we hypothesised that a microcolony surrounded by competitor microcolonies would have little impact on competitive outcome as growth would be ultimately limited. On the other hand, microcolonies located close to the edge of the biofilm would be free to expand radially and thus could make a more significant contribution to the competitive outcome (for an example timelapse video see Movie S3). Hence, we explored whether the competitive outcome was correlated to or even determined by the microcolonies’ *potential for radial expansion*. To do this requires the definition of an appropriate measure or score for this potential as follows. First, a circle is drawn that encloses the initial distribution of microcolonies. Second, each point on the circle is associated with the nearest microcolony and assigned to that strain. Third, the total arc length on the circle associated with each strain is computed. Finally, the *access to free space score* (AFS score) for strain *B*_1_, denoted AFS_1_, is then computed as the ratio of the total arc length associated with *B*_1_ to the circumference of the circle. It is straightforward to confirm that the access to free space score for strain *B*_2_, AFS_2_ = 1 — AFS_1_. See Section S4.2 and Figure S3 for a mathematically rigorous definition of the AFS score.

We explored the utility of the access to free space score using *N* = 6 and *N* = 824 as representatives of low and intermediate founder cell densities, respectively. We increased the number of model realisations to 5000 for each of the selected values of *N* to ensure improved accuracy of our data analysis. The AFS score was then calculated for each of the 10000 initial conditions (see examples Figure 3A and B). On completion of each of the simulations, the corresponding competitive outcome score was computed. Analysis of these data confirmed that access to free space accurately predicts competitive outcome: for each fixed founder density, the AFS score unfolds the variation shown in Figure 2D, yielding a positive, linear relationship between access to free space and competitive outcome (Figure 3C and 3D). For each of the selected values of *N*, initial configurations of microcolonies with a low AFS score predictably generated a low competitive outcome (score close to zero). Correspondingly, initial configurations with a high AFS score predictably generated a high competitive outcome (score close to 1). The slope of this linear relationship provides a deterministic measure of the range of competitive outcomes for a given founder density (cf. Figure 3C and 3D). The remaining small variation in the relationship is generated by the necessity to bin data into finite sets and by cases for which initial founder patch configurations have identical AFS scores but generate slightly different competitive outcomes. These latter cases occur because the AFS score does not account for dynamics within the confines of the footprint of the inoculum. For example, if microcolonies of one strain are placed so they initially *encircle* those of the other, then small changes to the positions of the encircled strain’s microcolonies will not change the AFS score. However, these *small* changes in initial configuration will cause *small* changes in competitive outcome. Moreover, due to the small size of the inoculum compared to the size of the mature biofilm, these small variations become negligible at larger simulation times.

**Figure 3:**
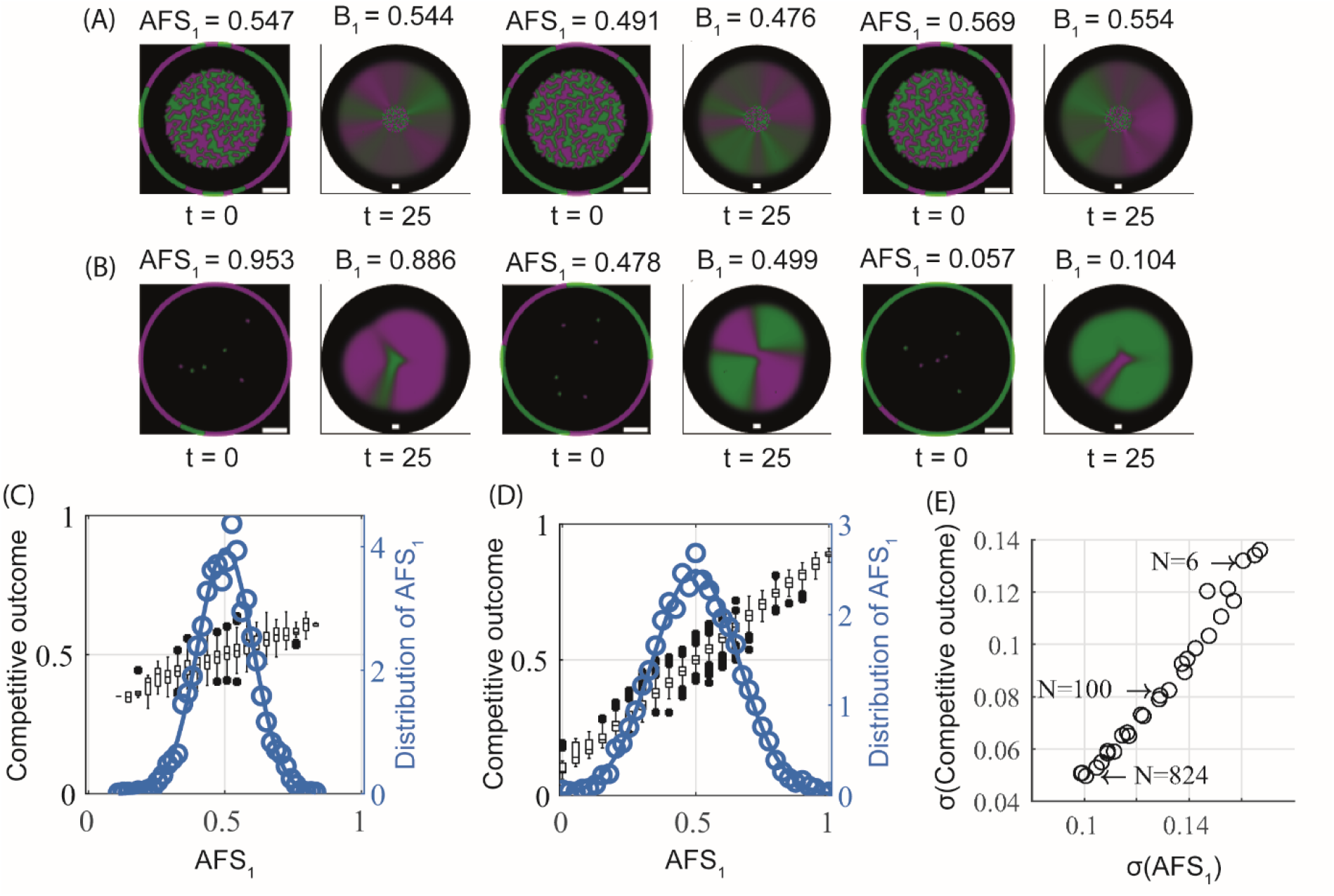
Access to free space determines competitive outcome. (A, B) Example model realisations for different founder densities. All plots show system’ initial conditions (*t* = 0) with the reference circle used to compute the access to free space score (the circle is rescaled for visualisation purposes) and outcomes after 25 time units. The founder densities are (A) *N* = 824 and (B) *N* = 6. Plots visualising the systems’ states at *t* = 0 show a blow-up of the subdomain *Ω*_0_; plots visualising outcomes at *t* = 25 show the full computational domain *Ω* (black background). The scalebars are seven unit lengths long. (C, D) The relation between the access to free space score *AFS*_1_, and competitive outcome is shown for (C) intermediate founder density (*N* = 824) and (D) low founder density (*N* = 6). Data are obtained from 5000 model realisations and cover the continuum of *AFS*_1_. The observed probability density function for AFS is shown (circular markers); along with the density function of a fitted normal distribution (*μ* ≈ 0.5, *σ* ≈ 0.10 in (C), *μ* ≈ 0.5, *σ* ≈ 0.16 in (D)) (solid line). (E) The relation between the standard deviations of the access to free space score *AFS*_1_ and the competitive outcome. Each data point (circle) represents a different founder density and contains information from 1000 model realisations

We subsequently established that the predictive power of the access to free space score was maintained across the range of founder densities considered in the model. Additionally the variation in the AFS score was shown to decrease with increasing founder density (cf. Figure 3C and D). Further, we revealed an approximately linear relationship between variation in AFS score and variation in competitive outcome (Figure 3E). Therefore, on increasing founder density, the observed decrease in variation in competitive outcome can be directly attributed to the decrease in variation in the AFS score.

### Dual strain single-isolate biofilm assays confirm modelling hypotheses

Next, we aimed to test the hypotheses put forward by the mathematical model. We selected an isogenic pair of *Bacillus subtilis* strains derived from isolate NCIB 3610 that constitutively expressed either the green fluorescent protein GFP (NRS6942, shown in green) or the blue fluorescent protein mTagBFP (NRS6932, shown in magenta). In line with the modelling assumption, the isolates were mixed in a 1:1 ratio at a defined initial cell density (we used an OD_600_ of 1) and this cell culture was serially diluted prior to inoculating the colony biofilms (Section S7). Thus, biofilms were inoculated using ~10^6^ CFUs and dilutions in 10-fold increments to order 1 CFU. For each founder density, 12 technical replicates were performed to provide a meaningful sample size, and the experiment was repeated on three independent occasions. We used a non-destructive colony biofilm image analysis approach, to measure the relative mass (and hence the *competitive outcome*) of the two isogenic strains at 24h, 48h, 72h after inoculation (see Section S10). We confirmed that the output from the image analysis correlated well with data generated by disruption of the colony biofilm and analysis of the relative strain proportions determined using single cells analysis by flow cytometry (Figure 4A) (see also (30)). The mTagBFP labelled strain consistently performed marginally worse than the GFP labelled competitor at high founder densities in co-culture, which suggests some impact on competitive fitness (Figure 4B and C).

**Figure 4:**
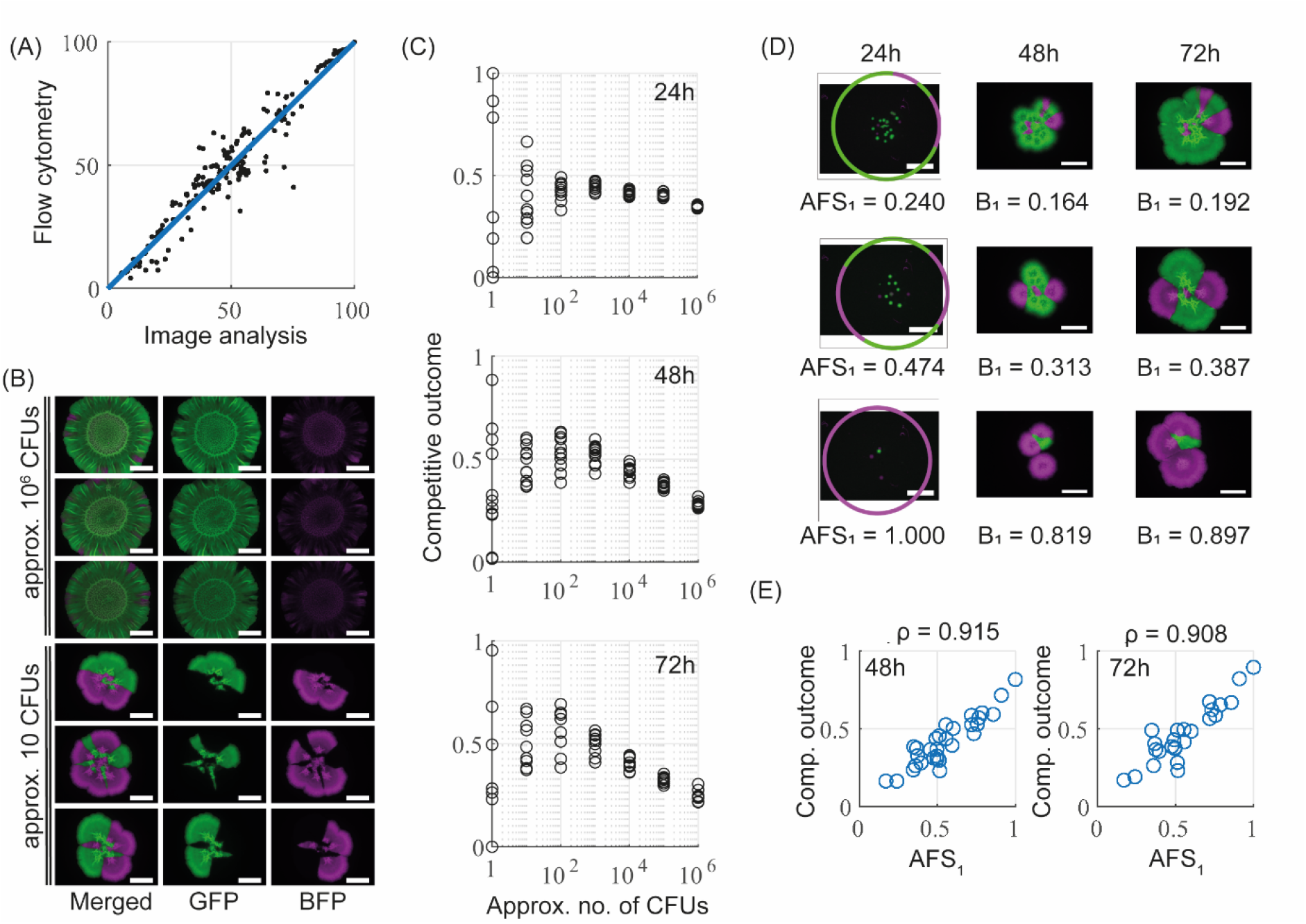
Experimental data confirm modelling hypotheses. (A) Comparison of image analysis with flow cytometry. A scatter plot comparing measurements of relative density of the mTagBFP-labelled strain obtained from image analysis and flow cytometry is shown. Each data point corresponds to one biofilm, which was imaged before being analysed by flow cytometry. The data contains measurements taken from all strain pairs, all founder densities, and all time points. The solid blue line shows the identity *x* = *y*, with the coefficient of determination being *R*^2^ = 0.91. (B) Example images of single-strain biofilms consisting of GFP (green) and mTagBFP (magenta) labelled copies of 3610. Taken after 72h of incubation and shown for two different founder densities (scalebar 5 mm). (C) Strain density data. Competitive outcome measurements taken after 24h, 48h and 72h of biofilm incubation. Plotted are technical repeats from one biological repeat of the experiment. The full data set is presented in Figure S4A. (D) Example visualisations of access to free space calculations. Three example biofilms images at d 24h (left), 48h (middle) and 72h (right). The strains are as described in (B). Images at 24h show the reference circle used for the access to free space score. (E) The relationship between access to free space and competitive outcome. AFS was calculated from images taken at 24h, and competitive outcome after 48h (left, *n* = 30) and 72h (right, *n* = 25). The linear correlation coefficient *ρ* is indicated.

Our experimental analysis proved consistent with the model output. High founder densities resulted in a broadly homogenous distribution of both strains over the footprint of the biofilm, while low founder densities led to a high degree of spatial segregation of the strains within the mature biofilm (Figure 4B, see also (14)). Additionally, analysis of experimental data confirmed that variability in the measured *competitive outcome* increased with decreasing founder density (Figure 4 B and C, Figure S4A). For founder densities equivalent to ~10^3^ to ~10^6^ CFUs, the competitive outcome was consistent across each set of technical replicates. By contrast, for founder densities of ~1 – 10^2^ CFUs, the competitive outcome was variable across each set of technical replicates. Moreover, variability in the competitive outcome score, at all initial founder densities, was marginally amplified over time.

We assumed the process of repeated dilution and selection of the inoculum volume may not guarantee an exact cell count and/or initial strain ratio of 1:1 at lower founder densities. Indeed, for low founder densities after 24 hrs incubation, we observed inconsistencies in the number and ratio of CFUs deposited (Figure S4B). We therefore considered whether this inherent experimental variability contributed to the observed variability in competitive outcome. To explore this in more detail, we first implemented a combinatorial argument that mathematically simulated the process of selecting the small inoculum volume from a larger cell culture (see methods). This process identified a threshold of ~10^2^ CFUs below which variability in cell number and/or strain ratio could measurably deviate from their intended values. Above this threshold, the combinatorial argument revealed limited deviation from the intended values (Figure S5A). Coupling these theoretical predictions with our experimental observations (Figure S4B), we therefore concluded that any observed variability in competitive outcome cannot be a consequence of a measurable deviation in the inoculum composition for colony biofilms founded with ~10^2^ CFUs or higher.

We next wanted to determine whether the predictive power of the access to free space score could be used to connect experimental initial configurations of the bacteria with the observed competitive outcome. To do this accurately, we required that the founding bacteria remained spatially separated as small colonies until an image was taken at 24 hrs (the earliest imaging time-point, see Figure 4D). This required using founder densities lower than 10^2^ CFUs. However, the computationally implied (and experimentally observed) inconsistencies in initial strain ratios and cell counts at these densities raised the question of whether access to free space could still accurately predict competitive outcome. To test this, we repeated our Monte Carlo simulations of (1) in which the number of initial microcolonies for each strain was drawn from an appropriate normal distribution, rather than being a fixed number and in a 1:1 ratio. Analysing the resulting model data confirmed that the predictive power of the access to free space score was robust to any “naturally-occurring” variation in the initial strain ratio (Figure S5B). Correspondingly, our analysis of the experimental data revealed a strong correlation between a strain’s access to free space score and the competitive outcome measured at 48h and 72h after incubation (Figure 4E).

### A modelling framework for non-isogenic strains

We have established that for isogenic strains, the initial configuration of founding bacteria determines the competitive outcome in a “race for space” and that the access to free space score can accurately predict which strain will dominate. A natural question that follows is what would happen if this race for space was influenced by other competitive mechanisms such as active killing mechanisms and/or different growth rates. Therefore, we considered the effect of introducing a (contact-dependent) killing mechanism by extending our theoretical framework (1). Moreover, constants describing the ratios between the strain’s maximum growth rates (*r*), diffusion rates (*d*) and killing coefficients (*c*) were introduced to allow for the possibility of differences in strain properties. This resulted in the following system obtained after a suitable nondimensionalisation (see supplementary information):

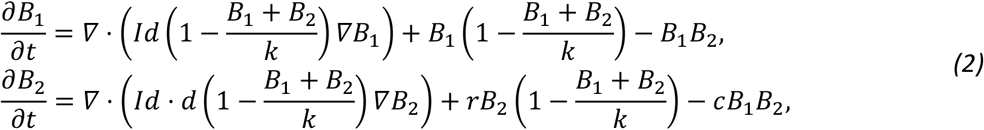

Here, the indicator function *Id* = 1 if *B*_1_ + *B*_2_ ≤*k* and Id = 0 otherwise. To start, strains were assumed to possess identical growth dynamics in the absence of competitors (i.e. r= 1, *d* = 1), but to significantly differ in their ability to kill the competitor strain. For the simulations we set *c* = 0.2 representing a five-fold difference in kill efficiency, with *B*_2_ being the more effective killer. A linear stability analysis of model (4) confirmed that in this case and for a homogeneous initial distribution of the strains in a 1:1 ratio, *B*_2_ wins the interaction. For this reason, we therefore refer to *B*_2_ as the (intrinsically) *stronger strain* and to *B*_1_ as the (intrinsically) *weaker strain* in the following.

The assumption of identical growth behaviour allowed us to focus on the impact on competitive outcome of antagonistic interactions. However, we anticipated that this assumption was unlikely to hold for non-isogenic strains in experimental settings and therefore we examined (as will be discussed later) the impact of changes to the parameters *r*, *d* and *c*. Subsequently, we showed the effect of such parameter variation to be limited.

### Spatial segregation induced by low founder densities enables coexistence

In the context of contact-dependent killing, low founder densities are expected to offer protection for the weaker strain by driving spatial segregation and the formation of enclaves. Test simulations supported this prediction. Model realisations with high (spatially uniform initial conditions) and intermediate (*N* = 824) founder densities consistently led to competitive exclusion of the weaker strain (Figure 5 A and B, Movies S4 and S5), while model realisations with low founder densities (*N* = 6) resulted in coexistence with the strains being spatially segregated (Figure 5C, Movie S6). As with the isogenic pairings, low founder densities generated significant variation in competitive outcome (Figure 5C). In particular, outcomes were observed for which the weaker strain *B*_1_ coexisted with, *and could even outperform*, the stronger strain *B*_2_. To better understand the impact of founder density, we performed Monte Carlo simulations with 1000 independent model realisations for each *N* in our test range. Data from these simulations revealed both the mean and variation of competitive outcome for the weaker strain increased with decreasing founder density (Figure 5D).

**Figure 5:**
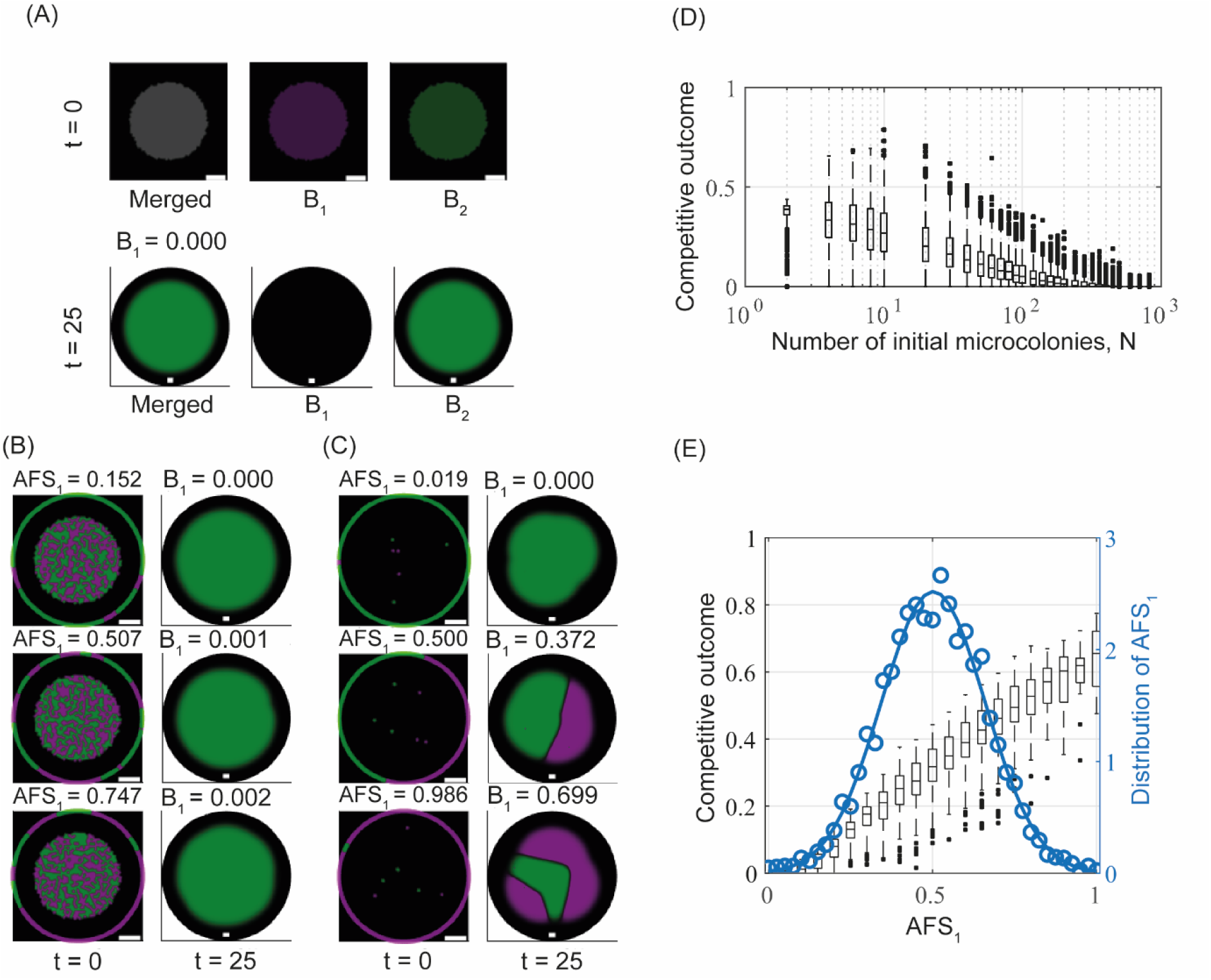
Modelling data for a non-isogenic strain pair. (A-C) Example model realisations for high (A), intermediate (B) and low (C) founder density are shown. In (A), the Merged image channel shows both strains (grey colour corresponds to overlap), the *B*_1_ and *B*_2_ channels only show single strain filters of the plot. In (B,C), only the Merged channel is shown. Plots visualising the systems’ states at *t* = 0 show a blow-up of the subdomain *Ω*_0_ and the circles used to calculate the access to free space scores around the initial conditions are not to scale. Plots visualising outcomes at *t* = 25 show the full computational domain *Ω* (black background). The scalebars are seven unit lengths long. (D) The relation between founder density and competitive outcome. Each boxplot contains data from 1000 model realisations. (E) The relation between the access to free space score *AFS*_1_, and competitive outcome for one fixed founder density (*N* = 6). Data are obtained from 5000 model realisations and covers the continuum of *AFS*_1_. The observed probability density function for AFS is shown (circular markers); the density function of a fitted normal distribution (*μ* ≈ 0.5, *σ* ≈ 0.16) as a solid line. The observed probability density function for AFS is shown (circular markers); the density function of a fitted normal distribution (*μ* ≈ 0.5, *σ* ≈ 0.16) as a solid line.

### Access to free space determines competitive outcome for low founder densities

The model consistently predicted competitive exclusion of the weaker strain at higher *N* values (Figure 5A and B). Hence, in these cases, the access to free space score no longer provided a meaningful predictor of competitive outcome. Rather, the model predicted the outcome to be dominated by the antagonistic interactions driven by contact dependent killing. However, as detailed above, low founder densities (*N* = 6) resulted in a highly variable competitive outcome and therefore, we explored if the access to free space score remained an accurate predictor in this case. The simulation data confirmed that for this fixed number *N*, the access to free space score remained capable of accurately unfolding the observed variation in competitive outcome (Figure 5E). Initial strain configurations with a low access to free space score predictably generated a low competitive outcome. The reciprocal was also maintained where initial strain configurations with a high access to free space score predictably generated high competitive outcome. As above, this relationship was found to be linear with the slope providing a measure of the determinist range of competitive outcomes for a given founder density. The relationship between access to free space and competitive outcome was again shown to be robust to natural variation in the initial strain ratio inherent in low founding cell densities (Figure S5C).

We note that our modelling approach revealed coexistence remained possible over a range of growth rates (less than a two-fold differences), diffusion coefficients (less than three-fold differences), and most surprisingly, *any* values of the killing coefficient (Section S6 and Figure S9A,B and C). In particular, we showed that a strain required extreme killing efficiency (*c* very large) in order to compensate for being slower in growth (*d*,*r* >1) (Figure S9D). Finally, the predictive power of the access to free space score was preserved over the parameter range tested (Figure S9E and F).

### Selection of a competition partner

To experimentally test our modelling hypotheses, we needed to identify a suitable partner for NCIB 3610. We chose a *Bacillus subtilis* strain called NRS6153 (hereafter 6153). This selection was made because *1*) 6153 is a genetically competent wild type strain with no known auxotrophies ((31)); *2*) in liquid culture conditions the generation times of the two strains are within ~1.5-fold of each other (Figure 6A); *3*) under biofilm conditions, single strain biofilms of both strains have footprint sizes that are within ~2-fold of each other (Figure 6B). *4*) across a broad range of founder densities, the competitive outcome of an isogenic pairing of 6153 isolates in a colony biofilm is broadly similar to that of an isogenic pairing of 3610 strains, albeit with more variability in the competitive outcome at the 72-hour time point for high founder densities (cf. Figure 4C (Fig. S4A) and Figure 6C (Fig. S6A)); 5) when a colony biofilm is founded at high density with marked strains of 3610 and 6153 starting at an initial 1:1 ratio, 6153 is consistently outcompeted by 3610 (and hence defines 3610 as the *stronger strain* in the context of this study) (Figure 6D); and 6) the mode of competition during co-culture in the colony biofilm is inferred as contact dependent as no obvious zone of growth inhibition was observed upon interrogation using an antibiosis halo formation assay (Figure 6E).

**Figure 6:**
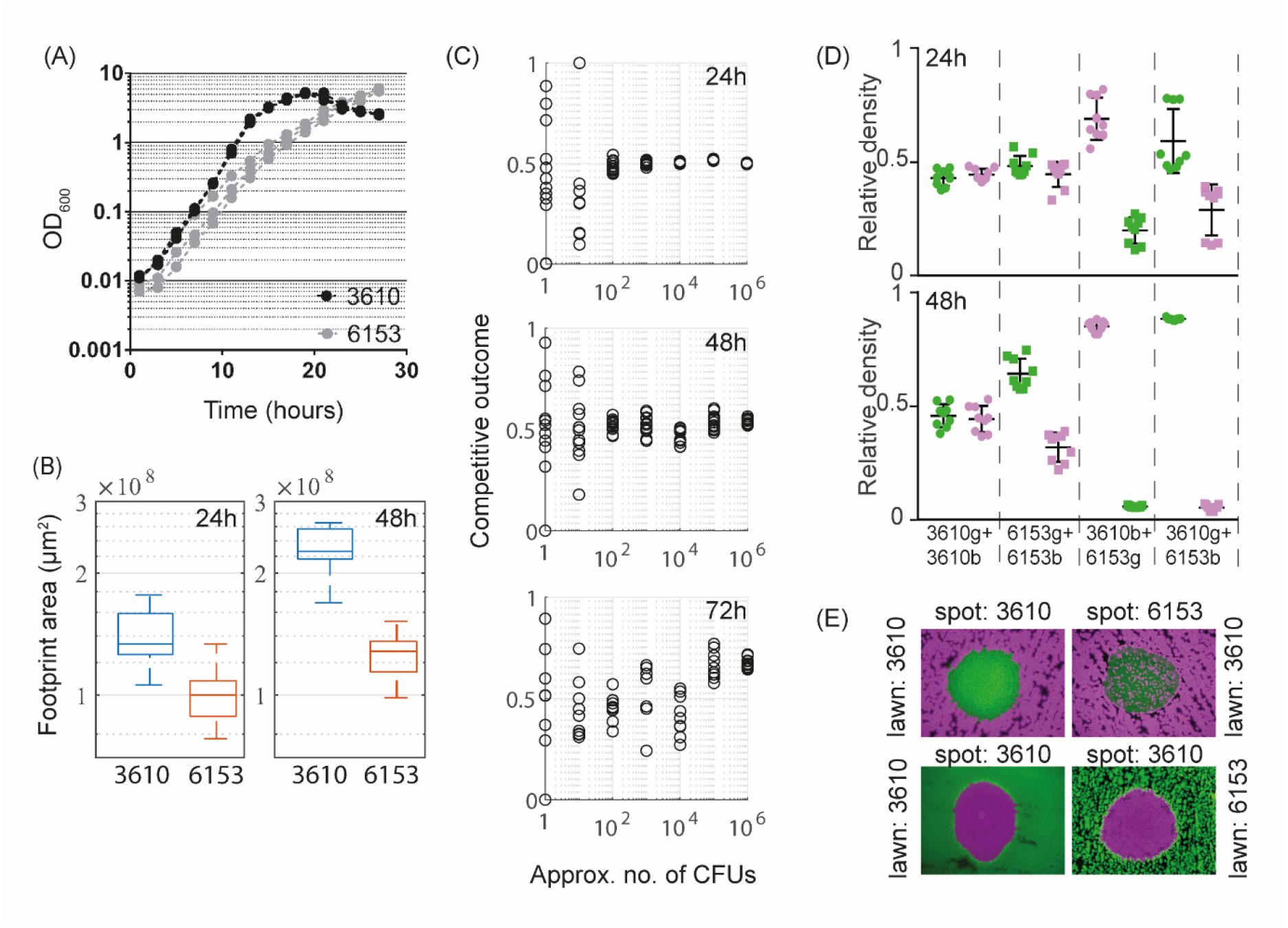
Selection of a competitive strain. (A) Growth curves of 3610 (black) and 6153 (grey) in MSgg cultures at 30°C. The three lines shown for each isolate represent separate biological repeats. (B) Biofilm footprint area of single-strain 3610 and 6153 biofilms. Data from 18 and 16 biofilms is shown for the 24h and 48h timepoint, respectively. (C) Competitive outcome data from colony biofilm assays of isogenic 6153 biofilms are shown after 24h, 48h and 72h of incubation. Plotted are the technical repeats from one biological repeat. The full data set is presented in Figure S6A. (D) Flow cytometry data of mixed biofilms grown for 24, 48 and 72 hours at 30°C on MSgg media. Isolate names followed by “g” represent strains constitutively expressing GFP, (green on the graph). Isolate names followed by “b” indicate strains constitutively expressing mTagBFP, (magenta on the graph). Three biological and three technical replicates were performed for each strain mix and timepoint and all data points are shown. The error bars represent the mean standard deviation. (E) Halo formation assays on MSgg agar plates at 24h of growth. Strains expressing mTagBFP (magenta) and GFP (green) are shown.

### Dual-isolate biofilm assays confirm modelling hypotheses

We performed dual strain biofilm assays competing 3610 and 6153 over a wide range of founder densities. These competitive assays confirmed the modelling hypothesis that coexistence within a non-isogenic strain pair is enabled by spatial segregation in biofilms inoculated at low founder densities (Figure 7A). Under such conditions, the intrinsically weaker strain (6153) formed spatial sectors and thus was able to coexist with the stronger strain (3610) through spatial segregation. (Fig. 8A and 8B). In contrast, and again as predicted by the model (and reported during the selection of strain 6153 as a competition partner), for biofilms inoculated at high founder density, the competitive outcome was realised as the competitive exclusion of 6153 (Figure 7A, 7B and S6B). Finally, a computation of access to free space scores based on images taken after 24h of incubation showed strong correlation between a strain’s access to free space and competitive outcome after both 48h and 72h of incubation for both 6153 alone and when in co-culture with 3610 (Figure S7 and Figure 7C).

**Figure 7:**
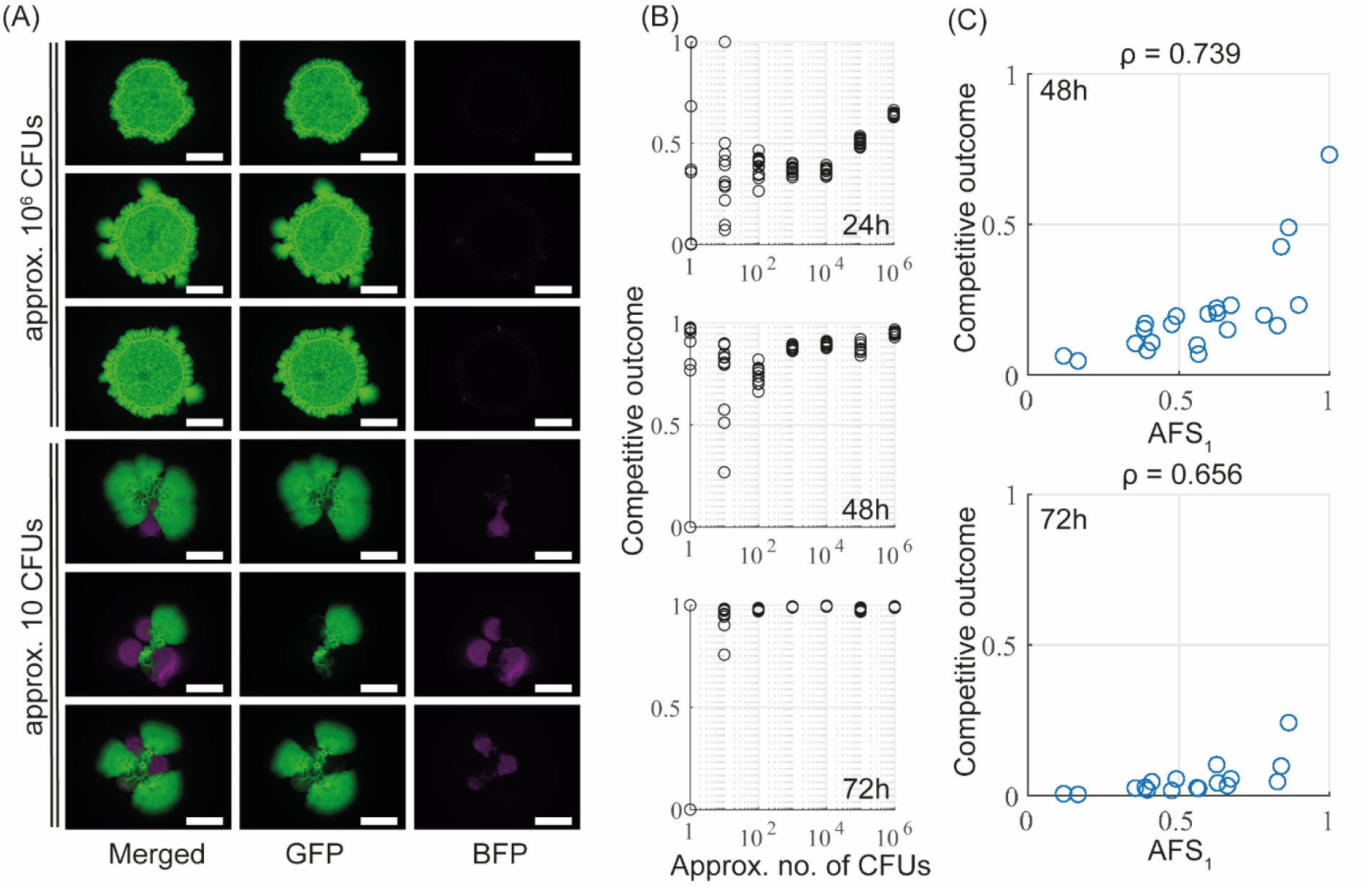
Experimental data for a non-isogenic strain pair. (A) Example dual-strain biofilms (3610 labelled with GFP (green), 6153 labelled with mTagBFP (magenta)). Images taken after 72h of incubation for two different founder densities. Scalebars as in Figure 2. (B) Competitive outcome data for 3610 in the 3610/6153 pair after 24h, 48h and 72h of biofilm incubation. Plotted are technical repeats from one biological repeat of the experiment. The full data set is presented in Figure S6B. (C) The relationship between access to free space of 6153, and competitive outcome. AFS was calculated based on images taken after 24h of biofilm incubation, and competitive outcome after 48h (top, *n* = 22) and 72h (bottom, *n* = 17).

## 3. Discussion

Significant advances in the understanding of competition dynamics within bacterial biofilms have been made in recent years (13, 14, 16–19, 26, 32). However, many basic questions remain open including how the initial configuration of the founding cells impacts the morphology and behaviour of the mature community. Here, using a multi-disciplinary approach, we have revealed that a *race-for-space* dominated competition dynamics in single- and dual-strain colony biofilms, even in cases where intra-species killing mechanisms strongly favoured survival of one strain over the other. In short, we established that range expansion dominated intra-species killing in biofilms inoculated with low founder densities. We showed that the resulting spatial structure supported coexistence of strains and revealed that it could lead to the counter-intuitive outcome of the *weaker* strain outperforming the *stronger* in terms of competitive outcome (relative biomass). Moreover, we established that a particular measure of the initial configuration of founder cells reliably predicted the resulting spatial structure and the relative densities of the strains in the mature biofilm. Our predictive measure, which characterises a strain’s *access to free space*, enabled us to disentangle experimentally observed variability in structure and competitive outcome in low founder density biofilms revealing this to be a natural, and predictable consequence of competition for space. This predictor proved to be remarkably robust to changes in the strain properties and initial strain ratio.

Our model system focussed on strain pairs that compete for space and interact through contact-dependent killing mechanisms. However, we hypothesise that results could be extended to strains interacting through contact-independent antagonistic actions, such as the secretion of diffusible antimicrobials (33). It is known that certain toxins have a restricted ability to penetrate cell colonies far from their production site (34). Hence, provided the spatial scale on which the contact-independent killing mechanism acts is sufficiently short, spatial segregation induced by the configuration of founder cells is still likely to drive dynamics very similar to those observed for contact-dependent mechanisms. In other words, although the *mechanisms* of contact-dependent and independent killing are different, the situation studied here can be viewed, theoretically in any case, as the behaviour in the limit as the toxin penetration depth tends to zero i.e. the toxin only has an impact very close to the producing cell. Tests of this hypothesis could be the subject of future work and may be performed through an extension of our theoretical framework through the explicit description of toxin dynamics and an experimental approach using a different choice of strains. In this way, it may be possible to determine critical penetration depths at which the competitive dynamics alter significantly from those discussed here.

Consistent with previous studies, we have shown that spatial segregation provides protection from contact-dependent antagonistic actions and is induced by low founder densities (14, 16, 17, 21, 26). Our findings highlight that the spatial structure induced by spatial segregation favours strains that would be outcompeted in unstructured environments and allows them to persist in colony biofilms. It is worth noting that low founder density is not the only mechanism that can induce spatial segregation of cell linages in a biofilm. Spatial structure can also be induced by genetic drift due to the small size of the population within the biofilm edge that contributes to radial expansion (13, 18, 19, 35). However, spatial segregation via genetic drift is a gradual process that requires strains to coexist without spatial structure in the biofilm centre (13, 18, 19, 36). It is thus unlikely to affect biofilm phenotype if antagonistic interactions that prevent coexistence in the biofilm centre dominate competitive interactions. Indeed, spatial structure induced by genetic drift is commonly associated with prevention of exploitation of co-operators by cheaters in social dynamics, rather than protection from antagonistic actions (13, 35, 37, 38).

The development of a deeper understanding of competition dynamics in multi-strain biofilms is an essential precursor for the optimal design and implementation of industrial applications. For example, biofilm-forming microbial species in the rhizosphere can mutualistically interact with plant roots and are therefore used as biofertilizers and biopesticides. Biofilms on plant roots can supply plants with fixed nitrogen and provide protection from plant pathogens (39–41) in exchange for root exudates (42, 43), such as carbon (44). *B. subtilis* is a species widely used in biocontrol (40, 45, 46). However, *B. subtilis* has a large and open pangenome (47, 48) and only select isolates have been shown to possess traits associated with successful applications as biocontrol agents (49). A biocontrol agent can only be successfully applied if it manages to coexist (or outcompete) other strains already present. These and many other examples, such as the applications of biofilms in wastewater treatment (6), microbial fuel cells (50) and corrosion prevention (51), illustrate the need to better understand interstrain interactions within biofilms growing in complex and potentially continually changing environments. Therefore, we argue that future work focusing on enhancing our understanding of spatial structure and competition within multi-strain biofilms will be critical to optimise our ability to maximise the positive impact of biofilms.

## Acknowledgements

Work in the NSW, FAD and CEM laboratories is funded by the Biotechnology and Biological Science Research Council (BBSRC) [BB/P001335/1, BB/R012415/1]. MK is supported by a Biotechnology and Biological Sciences Research Council studentship (BB/M010996/1). We are grateful to Dr Natalie Bamford and other members of the Stanley-Wall lab for helpful discussions. We would like to acknowledge the Flow Cytometry and Cell Sorting Facility at the University of Dundee

## Data availability statement

Computational code is available on Github and has been archived by Zenodo (52). The experimental datasets have been achieved using BioStudies (53).

## Author Contributions using CRediT

Conceptualisation: LE, NSW, FAD

Data curation: LE, MK

Formal Analysis: LE, MK

Funding acquisition: CEM, NSW, FAD

Investigation: LE, MK

Methodology: LE, MK, GB, FAD

Project administration: NSW

Resources: MK

Software: LE, GB

Supervision: CEM, NSW, FAD

Validation: LE, MK, NSW, FAD

Visualization: LE, MK

Writing – original draft: LE, NSW, MK, FAD

Writing – review & editing: LE, MK, GB, CEM, NSW, FAD

