## Supplementary material for "Founder cell configuration drives competitive outcome within colony biofilms"

Lukas Eigentler[1,2], Margarita Kalamara[1], Graeme Ball[3], Cait E. MacPhee[4], Nicola R. Stanley-Wall[1,*], Fordyce A. Davidson[2,*]

[1] Division of Molecular Microbiology, School of Life Sciences, University of Dundee, Dundee DD1 5EH, United Kingdom
[2] Division of Mathematics, School of Science and Engineering, University of Dundee, Dundee DD1 4HN, United Kingdom
[3] Dundee Imaging Facility, School of Life Sciences, University of Dundee, Dundee DD1 5HN, United Kingdom
[4] School of Physics and Astronomy, The University of Edinburgh, Edinburgh EH9 3FD, UK

### S1 Supplemental figures

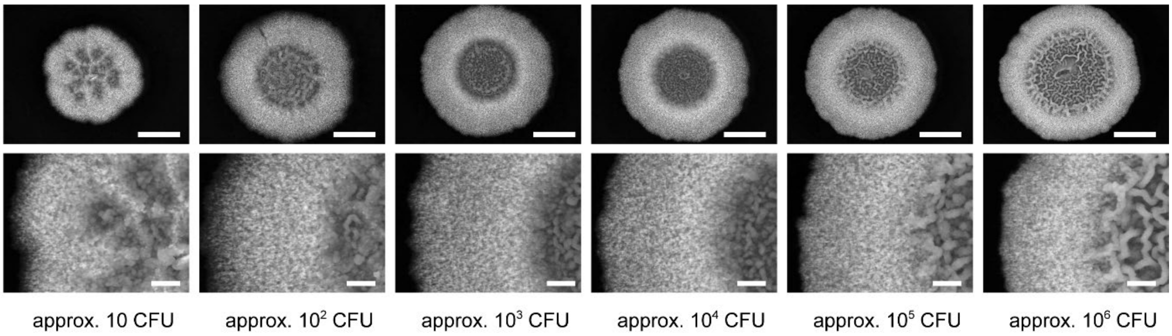

**Figure S*1*: The mature biofilm morphology is robust to the number of founding cells.** B. subtilis strain NCIB 3610 was used to establish colony biofilms using a range of founder densities. The biofilms were grown at 30°C for 48 hours prior to photography. The scale bar represents 5 mm.

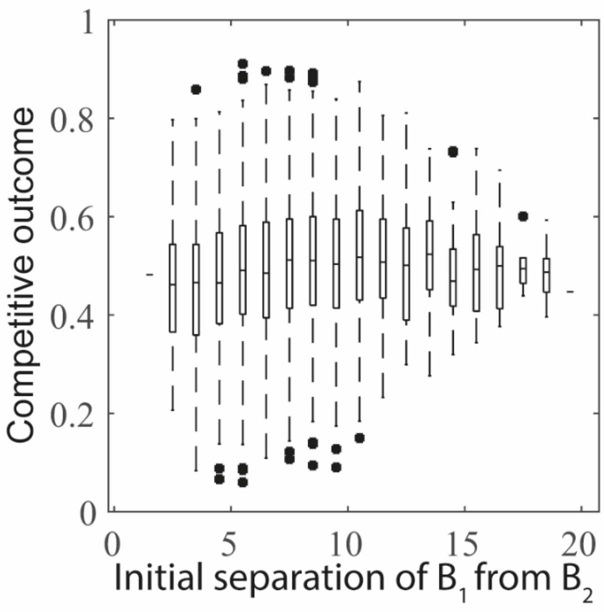

Figure S2: **Initial interstrain separation cannot predict competitive outcome in isogenic strain pairs.** The mean initial separation of $B_1$ microcolonies to their nearest $B_2$ microcolony is plotted against the corresponding competitive outcome. The initial separation between the microcolonies was split into bins for visualisation purposes. No clear relation between the two quantities exists.

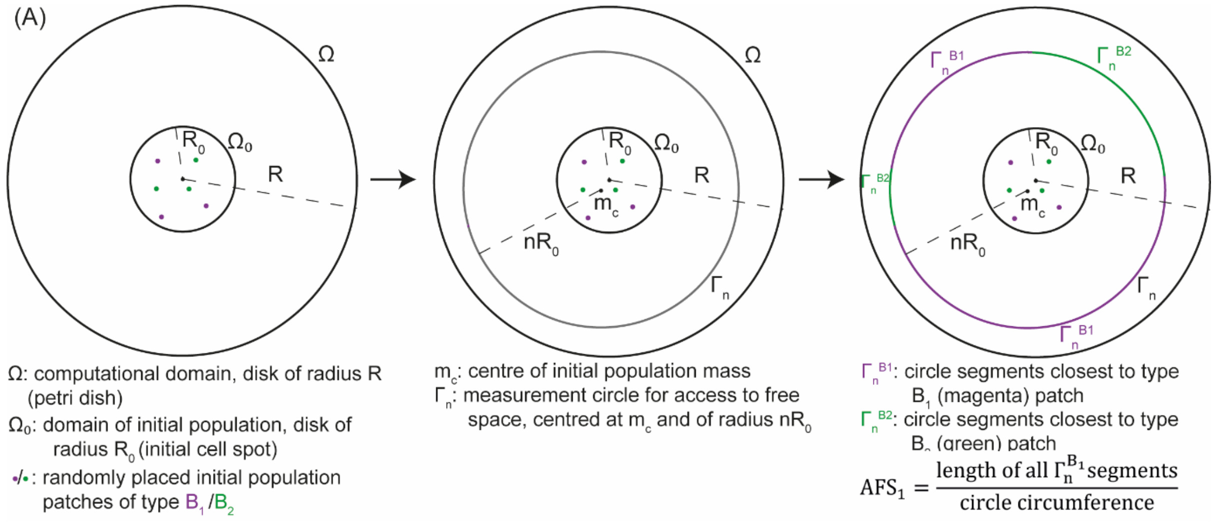

*Figure S3: (A)* ***Schematic of the derivation of the access to free space score.*** *A reference circle $\Gamma_n$ of radius $nR_0$ is drawn centred at $m_c$, which is the centre of mass of the initial population. Circle segments are assigned to set $\Gamma_n^{B_1}$if the closest strain patch belongs to $B_1$, and to $\Gamma_n^{B_2}$ otherwise. For ease of visualisation, these segments are given the same colour as the nearest patch. The colour of the circle changes at points where nearest patches of different strains are equidistant to the point on the circle. The ratio of the total length of the magenta segments to the circumference of $\Gamma_n$ defines the access to free space score $AFS_1$ of strain $B_1$, and correspondingly for strain $B_2$ (green).*

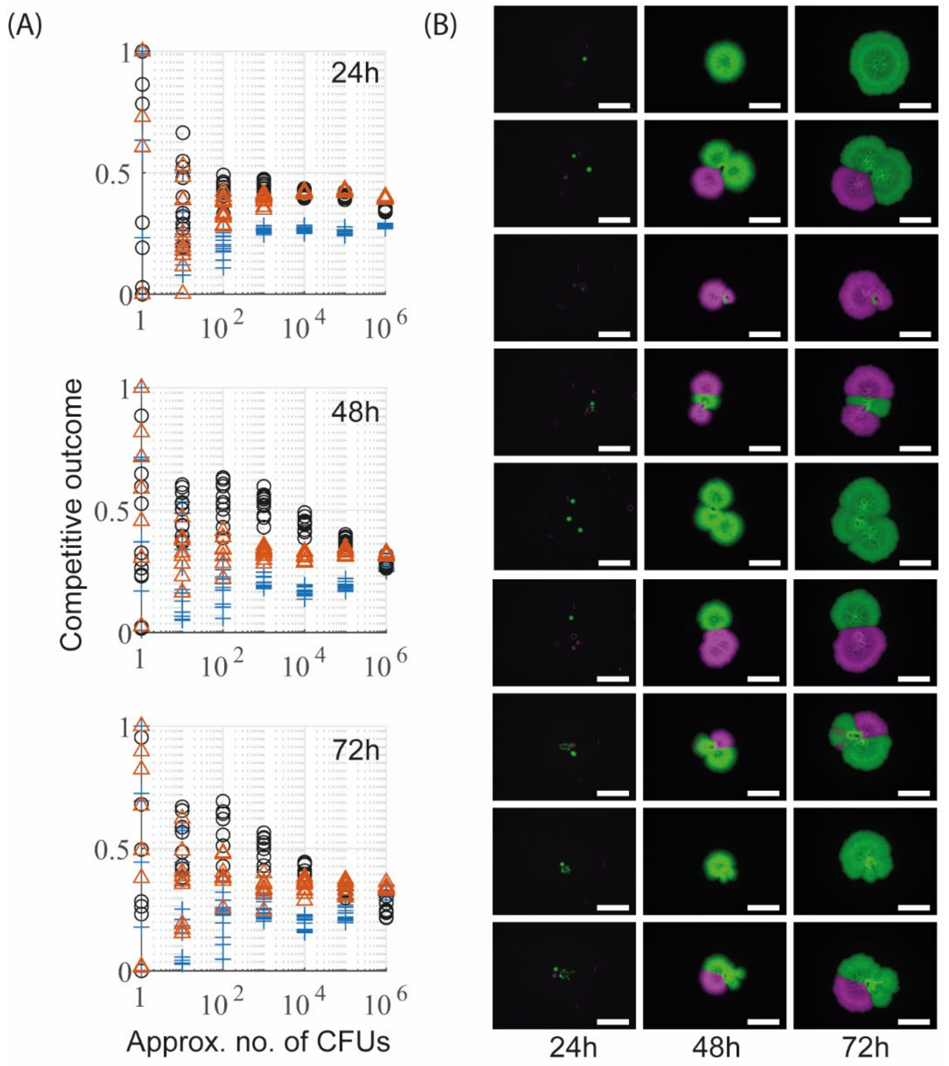

*Figure S4: (A)* ***Experimental data of all three biological replicates****. Competitive outcome data for one strain only from all three biological replicates are shown for the isogenic 3610 assay at 24h, 48h and 72h after biofilm inoculation. The colour and marker shape indicate data from the same biological replicate. For any given data point, the competitive outcome of the competitor strain is the given score subtracted from unity. (B) Variability at low founder density caused by inconsistencies in initial strain ratio. Images taken after 24, 48 and 75 hours of incubation of all technical repeats within one biological repeat of our 3610 isogenic strain pair assay, inoculated with founder densities of order 1 CFU.*

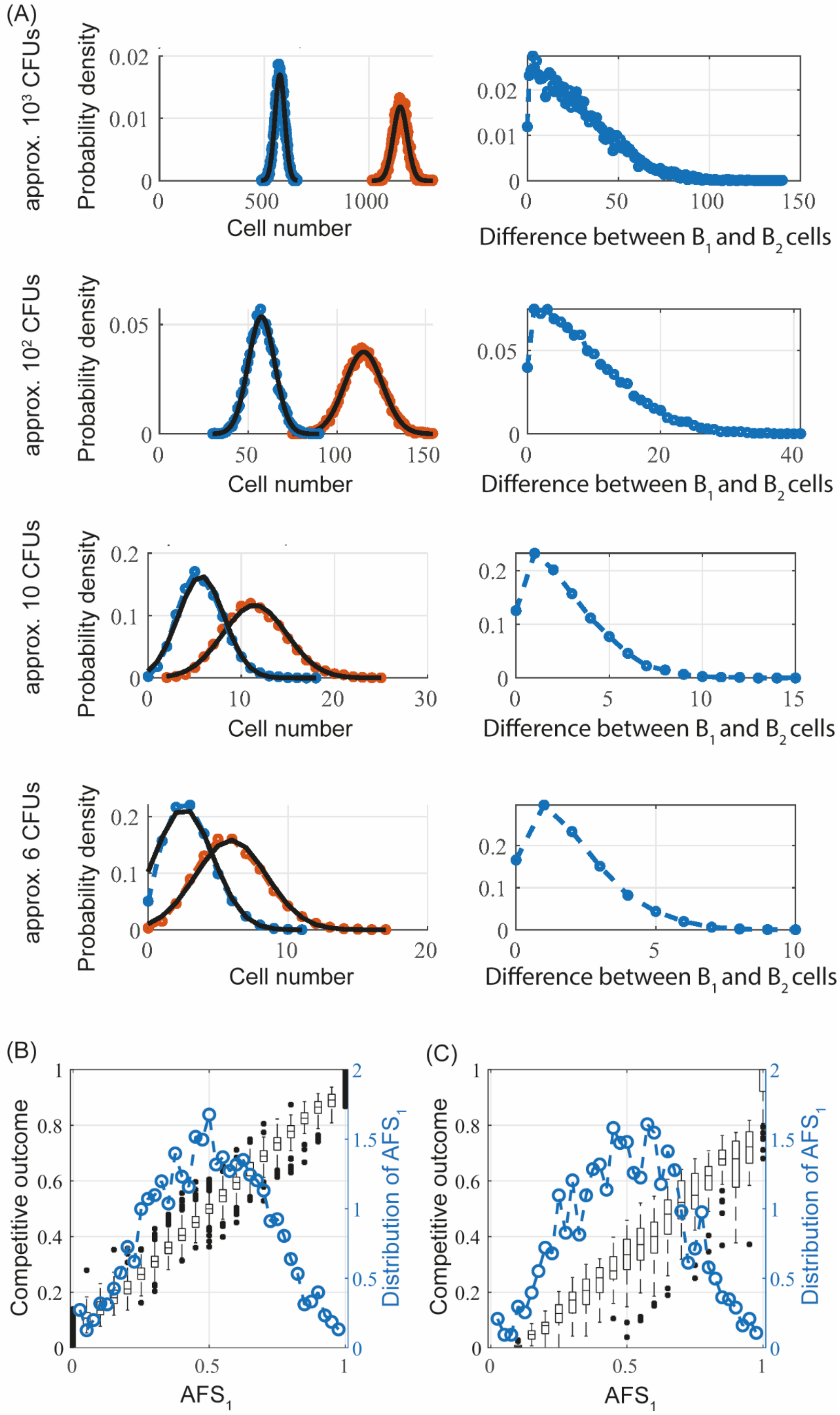

(A)
approx. $10^3$ CFUs
approx. $10^2$ CFUs
approx. 10 CFUs
approx. 6 CFUs
Probability density
Cell number
Difference between $B_1$ and $B_2$ cells
(B)
(C)
Competitive outcome
Distribution of $AFS_1$
$AFS_1$

*Figure S5:* ***Data from combinatorial cell picking model.*** *(A) Cell number distributions at different founder densities. Data obtained from a Monte Carlo simulation with 10000 replicates of our combinatorial cell picking model, representing the process of inoculating biofilms with* $10^3$*,* $10^2$*,* $10$ *and* $6$ *CFUs. The left column shows total cell numbers (red) and cell numbers of the strain labelled as* $B_1$ *(blue). The distribution of the second strain is the same as that for* $B_1$*. Both distributions are approximately normal, as indicated by the fitted (truncated) normal distributions (solid black curve). The right column shows the distribution of the difference between cell numbers of both strains. (B,C) Access to free space determines competitive outcome independent of initial strain ratio. The relation between the access to free space score* $AFS_1$ *and competitive outcome is shown for an isogenic strain pair (B) and a non-isogenic strain pair (C). The data are obtained from a Monte Carlo simulation (5000 realisations each), in which initial cell numbers are picked at random from the distribution calculated by our combinatorial model representing a founder density equivalent to approx. 6 CFUs to allow for comparisons with results presented in the main text. Circular makers connected by dashed lines visualise the observed distribution of* $AFS_1$*.*

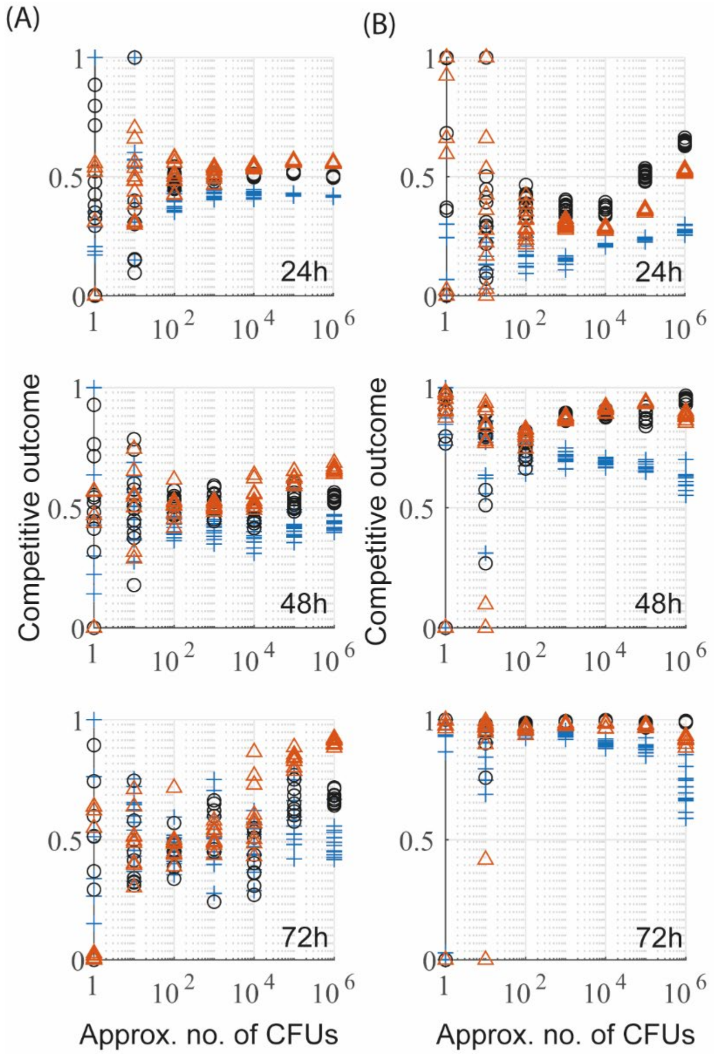

*Figure S6:* ***Experimental data of all three biological replicates.*** *Competitive outcome data of one strain only from all three biological replicates are shown for isogenic 6153 pair (A) and the competitive 3610/6153 pair (B) assays at 24h, 48h and 72h after biofilm inoculation. The colour and marker shape indicate data from the same biological replicate.*

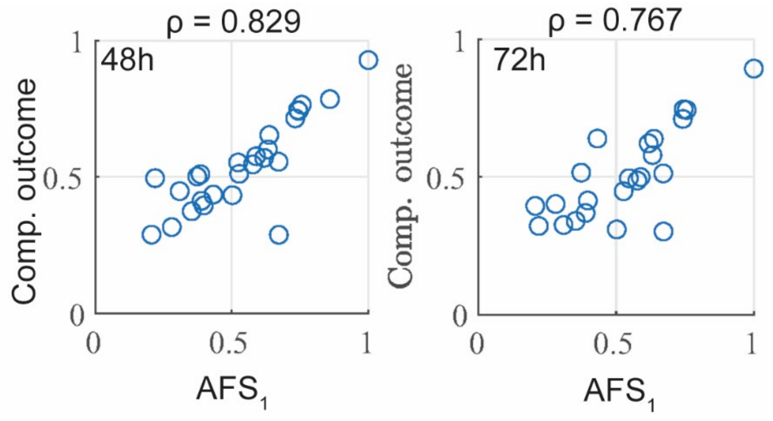

*Figure S7:* ***Access to free space as a predictor of competitive outcome for the isogenic 6153 pair.*** *The relation between access to free space, calculated based on images taken after 24h of biofilm incubation, and strain density after 48h (left,* $n = 26$*) and 72h (right,* $n = 23$*) is shown.*

### S2 Supplemental movies

Movie S1: “Movie_S1_high_founder_dens_isogenic.avi”
Timelapse video of the mathematical model for isogenic strains, initialised used spatially homogeneous conditions, representing high founder density.

Movie S2: “Movie_S2_intermediate_founder_dens_isogenic.avi”
Timelapse video of one example realisation of the mathematical model for isogenic strains, initialised using $N = 824$ microcolonies, representing intermediate founder density.

Movie S3: “Movie_S3_low_founder_dens_isogenic.avi”
Timelapse video of one example realisation of the mathematical model for isogenic strains, initialised using $N = 6$ microcolonies, representing low founder density.

Movie S4: “Movie_S4_high_founder_dens_nonisogenic.avi”:
Timelapse video of the mathematical model for non-isogenic strains, initialised used spatially homogeneous conditions, representing high founder density.

Movie S5: “Movie_S5_intermediate_founder_dens_nonisogenic.avi”
Timelapse video of one example realisation of the mathematical model for non-isogenic strains, initialised using $N = 824$ microcolonies, representing intermediate founder density.

Movie S6: “Movie_S6_low_founder_dens_nonisogenic.avi”
Timelapse video of one example realisation of the mathematical model for non-isogenic strains, initialised using $N = 6$ microcolonies, representing low founder density.

#### S3 Models and nondimensionalisations

The mathematical model (1) in the main text is nondimensionalised. In dimensional form, it is

$$\frac{\partial \tilde{B}_1}{\partial \tilde{t}} = \nabla \cdot \left( Id \cdot \tilde{d} \left(1 - \frac{\tilde{B}_1 + \tilde{B}_2}{\tilde{k}_{cap}}\right) \nabla \tilde{B}_1 \right) + \tilde{k}_{max} \left(1 - \frac{\tilde{B}_1 + \tilde{B}_2}{\tilde{k}_{cap}}\right),$$
$$\frac{\partial \tilde{B}_2}{\partial \tilde{t}} = \nabla \cdot \left( Id \cdot \tilde{d} \left(1 - \frac{\tilde{B}_1 + \tilde{B}_2}{\tilde{k}_{cap}}\right) \nabla \tilde{B}_2 \right) + \tilde{k}_{max} \left(1 - \frac{\tilde{B}_1 + \tilde{B}_2}{\tilde{k}_{cap}}\right). \qquad (S1)$$

The variables $\tilde{B}_1(\tilde{\boldsymbol{x}}, \tilde{t})$ and $\tilde{B}_2(\tilde{\boldsymbol{x}}, \tilde{t})$ (both units: $gm^{-2}$) denote the densities of two bacterial strains at time $\tilde{t} > 0$ ($s$) and spatial position $\tilde{\boldsymbol{x}} \in \Omega$, ($m^2$) where $\Omega \subset \mathbb{R}^2$ is the two-dimensional spatial domain. The parameter $\tilde{k}_{max}$ ($s^{-1}$) is the maximum growth rate coefficient of the isogenic strains $\tilde{B}_1$ and $\tilde{B}_2$. The carrying capacity $\tilde{k}_{cap}$ ($gm^{-2}$) denotes the maximum, sustainable population density. Finally, $\tilde{d}$ ($m^2 s^{-1}$) is the maximum diffusion coefficient of the strains and the indicator function $Id = 1$ if $\tilde{B}_1 + \tilde{B}_2 \leq \tilde{k}_{cap}$ and $Id = 0$ otherwise. A suitable nondimensionalisation is

$$\tilde{B}_1 = \tilde{k}_{cap} B_1, \qquad \tilde{B}_2 = \tilde{k}_{cap} B_2, \qquad \tilde{t} = \frac{1}{\tilde{k}_{max}} t, \qquad \tilde{\boldsymbol{x}} = \sqrt{\frac{\tilde{d}}{\tilde{k}_{max}}} \boldsymbol{x}. \qquad (S2)$$

Substitution of (S2) into (S1) yields the nondimensional model (1) in the main text.

The mathematical model (4) in the main text is nondimensionalised. In dimensional form, it is

$$\frac{\partial \tilde{B}_1}{\partial \tilde{t}} = \nabla \cdot \left( Id \cdot \tilde{d}_{\tilde{B}_1} \left(1 - \frac{\tilde{B}_1 + \tilde{B}_2}{\tilde{k}_{cap}}\right) \nabla \tilde{B}_1 \right) + \tilde{k}_{max,\tilde{B}_1} \tilde{B}_1 \left(1 - \frac{\tilde{B}_1 + \tilde{B}_2}{\tilde{k}_{cap}}\right) - \tilde{c}_{12} \tilde{B}_1 \tilde{B}_2,$$
$$\frac{\partial \tilde{B}_2}{\partial \tilde{t}} = \nabla \cdot \left( Id \cdot \tilde{d}_{\tilde{B}_2} \left(1 - \frac{\tilde{B}_1 + \tilde{B}_2}{\tilde{k}_{cap}}\right) \nabla \tilde{B}_2 \right) + \tilde{k}_{max,\tilde{B}_2} \tilde{B}_2 \left(1 - \frac{\tilde{B}_1 + \tilde{B}_2}{\tilde{k}_{cap}}\right) - \tilde{c}_{21} \tilde{B}_1 \tilde{B}_2. \qquad (S3)$$

As above, the variables $\tilde{B}_1(\tilde{\boldsymbol{x}}, \tilde{t})$ and $\tilde{B}_2(\tilde{\boldsymbol{x}}, \tilde{t})$ (both units: $gm^{-2}$) denote the densities of two bacterial strains at time $\tilde{t} > 0$ ($s$) and spatial position $\tilde{\boldsymbol{x}} \in \Omega$, ($m^2$) where $\Omega \subset \mathbb{R}^2$ is the two-dimensional spatial domain. The parameters $\tilde{k}_{max,\tilde{B}_1}$ and $\tilde{k}_{max,\tilde{B}_2}$ (both $s^{-1}$) are the maximum growth rate coefficients of $B_1$ and $B_2$, respectively. The carrying capacity $\tilde{k}_{cap}$ ($gm^{-2}$) denote the maximum, sustainable population density. The parameter $\tilde{c}_{12}$ ($m^2 g^{-1} s^{-1}$) denotes the amount of biomass of $B_1$ killed per second per unit biomass of strain $B_2$, and *vice versa* for $\tilde{c}_{21}$ ($m^2 g^{-1} s^{-1}$). Finally, $\tilde{d}_{\tilde{B}_1}$ and $\tilde{d}_{\tilde{B}_2}$ (both $m^2 s^{-1}$) are the maximum diffusion coefficients of $\widetilde{B_1}$ and $\widetilde{B_2}$, respectively. A suitable nondimensionalisation is

$$\tilde{B}_1 = \frac{\tilde{k}_{max,\tilde{B}_1}}{\tilde{c}_{12}} B_1, \qquad \tilde{B}_2 = \frac{\tilde{k}_{max,\tilde{B}_1}}{\tilde{c}_{12}} B_2, \qquad \tilde{t} = \frac{1}{\tilde{k}_{max,\tilde{B}_1}} t, \qquad \tilde{\boldsymbol{x}} = \sqrt{\frac{\tilde{d}_{\tilde{B}_1}}{\tilde{k}_{max,\tilde{B}_1}}} \boldsymbol{x},$$
$$k_{cap} = \frac{\tilde{k}_{cap} \tilde{c}_{12}}{\tilde{k}_{max,\tilde{B}_1}}, \qquad d = \frac{\tilde{d}_{\tilde{B}_2}}{\tilde{d}_{\tilde{B}_1}}, \qquad k_{max} = \frac{\tilde{k}_{max,\tilde{B}_2}}{\tilde{k}_{max,\tilde{B}_1}}, \qquad c = \frac{\tilde{c}_{21}}{\tilde{c}_{12}} \qquad (S4)$$

Substitution of (S4) into (S3) yields the nondimensional model (4) in the main text.

### S4 Methods for model analysis

#### S4.1 Model implementation

The mathematical models were solved numerically for 25 nondimensional time units using Matlab's PDE Toolbox (Version 3.5 (Release 2020b))(1), which implements a finite element method. We chose a triangulated mesh of linear geometric order, with all mesh elements being of approximately the same size. We performed checks to ensure that the choice of mesh size did not affect results presented. The spatial domain was chosen to be large enough so that populations could not reach the boundary within the time considered in the numerical simulations. Thus, boundary conditions (the derivatives of $B_1$ and $B_2$ in the direction of the outward normal are set to zero) did not have an impact on the solution dynamics. Unless otherwise stated, parameter values were $R = 10\sqrt{50}, R_0 = 0.2R$ (both models) and $d = r = 1, c = 0.2, k = 10$ (in (4) only). The target side length of mesh elements was set to $0.125\sqrt{50}$, except in the simulations for Fig. 6D, which required a finer mesh (target side length was halved) due to the large killing coefficients.

For both models, Monte Carlo simulations were performed for selected values of the number of initial microcolonies, $N$, with $N$ drawn from the set $S$ with

$$S \coloneqq \{2,4,6,8,10,20,30,40,50,60,70,80,90,100,120,140,160,180,200,240,280,320,$$

$$360,400,450,500,600,700,824\}.$$

#### S4.2 Definition of access to free space score

The access to free space scores $AFS_i, i = 1{,}2$, are defined as follows. First, for a given initial condition (initial configuration), we denote by $\mathcal{B}_i = \{\boldsymbol{x} \in \Omega: B_i(\boldsymbol{x}, 0) > 0\}$ the set of loci that are assigned to microcolonies of strain $B_i$. To define $\mathrm{AFS_i}$, we considered a circle $\Gamma_n = \{\boldsymbol{x} \in \mathbb{R}^2: \|\boldsymbol{x} - \boldsymbol{m_c}\| = nR_0\}$, where $n$ is a positive real number and $\boldsymbol{m_c}$ is the centre of mass of the whole initial population (Figure 3A). The number $n$ was chosen to be sufficiently large so that $\Gamma_n$ encompasses the whole of $\Omega_0$ (i.e. $n \geq 1$). For each point $\boldsymbol{x}_c$ on the circle we (i) determined the closest microcolony, (ii) recorded which strain occupied that microcolony and (iii) assigned the point $\boldsymbol{x}_c$ to the set $\Gamma_n^{\mathcal{B}_1}$ if the closest patch belongs to $\mathcal{B}_1$ and to $\Gamma_n^{\mathcal{B}_2}$ otherwise (Fig. S3). That is

$$\Gamma_n^{\mathcal{B}_i} = \{\boldsymbol{x_c} \in \Gamma_n: \min_{y \in \mathcal{B}_i} \|\boldsymbol{x_c} - \boldsymbol{y}\| < \min_{y \in \mathcal{B}_j} \|\boldsymbol{x_c} - \boldsymbol{y}\|\}. \qquad (5)$$

The *access to free space score* of $B_i$, $\mathrm{AFS_i}$, was then defined to be the ratio of the total length of circle segments in $\Gamma_n^{\mathcal{B}_i}$, denoted by $\ell(\Gamma_n^{\mathcal{B}_1})$, to the circumference of the circle $\Gamma_n$ i.e.

$$AFS_1 = \frac{\ell(\Gamma_n^{\mathcal{B}_1})}{2nR_0\pi}, \qquad AFS_1 = 1 - AFS_2. \qquad (6)$$

It is clear from its definition that the access to free space score depends on the choice of $n$, i.e. $\mathrm{AFS_i} = \mathrm{AFS_i}(n)$. It was established that even $n$=1 provided a reasonably accurate measure and increasing $n$ beyond 2 did not induce a significant change in $\mathrm{AFS_i}(n)$ (Section S4.3 and Figure S8). For consistency, we used $n = 5$ in all the analysis of our modelling data, and for brevity denoted $\mathrm{AFS_i}(5)$ by $\mathrm{AFS_i}$.

#### S4.3 Choice of R in access to free space score

The access to free space score $\mathrm{AFS_i}$ depends on the measurement radius $R \coloneqq nR_0$. A comparison of $\mathrm{AFS_i}$ with different values of $n$ highlights that no significant change to $\mathrm{AFS_i}$ occurs if $n > 2$ (Fig. S8D). This is highlighted by the decrease of the "width of the band" of densities $B_1$ predicted to occur for a given $\mathrm{AFS_i}$ with increasing $n$ (Fig. S8 A-C).

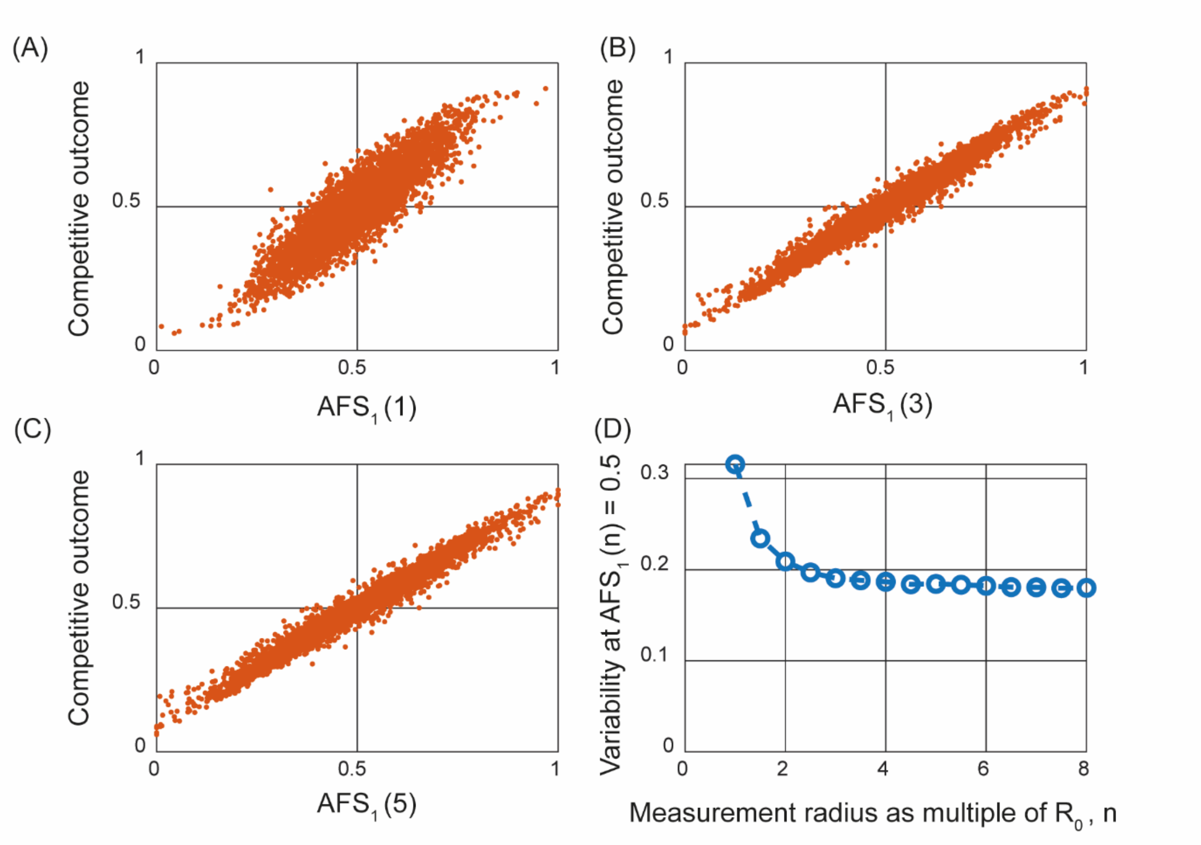

*Figure S8:* ***The impact of $n$ on the access to free space score.*** *(A-C) Scatter plots of outcome strain density $B_1$ against the access to free space score $AFS_1$ of the system's initial condition is shown for three different values of $n$. (D) Range of $B_1$ in which 95% of data with an access to free space score $AFS_1 \approx 0.5$ is contained. Significant changes to $AFS_1$ occur as $n$ increases from its minimum $n = 1$, but for sufficiently large $n$, changes in $n$ have negligible impact on $AFS_1$. Each data point represents one model realisations with $N = 6$ initial cell patches of two isogenic theoretical strains.*

#### S5 No variability in competitive outcome for initial conditions with two founder cells only

The case of $N = 2$ which each strain only occupies one single cell patch initially is special. This initial condition induces symmetry in the system around the axis that is perpendicular to, and bisects the line connecting the initial patches. This symmetry results in identical behaviour of both strains yielding a competitive outcome of $\approx 0.5$ consistently. Note the slight deviation from 0.5 which occurs due to the discretisation of the domain. It is noteworthy that this does not contradict our characterisation of competitive outcome by access to free space, discussed in the main text. If there are only two occupied cell patches initially, the system's centre of mass is the bisection point of the line connecting both patches. Hence the intersection points between the reference circle $\Gamma_n$ used to define the access to free space score $\mathrm{AFS_i}$ and the system's symmetry axis are equidistant to both cell patches and split $\Gamma_n$ into two segments of the same length ($\mathrm{AFS_1} = 0.5$).

#### S6 Model hypotheses are robust to differences in strain growth dynamics

Next, we considered the effect of allowing the growth, diffusion and killing parameters of the two strains to differ. A full sensitivity analysis that encompassed the whole parameter space for each founder density and corresponding large set of initial data was infeasible due to the high computational cost. Instead, for each founder density, we fixed an initial configuration that resulted in a typical competitive outcome (i.e. one that lied within the interquartile range given by our Monte Carlo simulations in the sense of Figure 2D). We then investigated the impact of changes to strain growth rate, killing coefficient and the diffusion coefficient.

Unsurprisingly our analysis revealed that increasing a strain's growth rate, diffusion coefficient, or killing efficiency led to an increased competitive outcome for that strain (Figure S9A,B and C). In particular, when the growth rate parameter $r \gtrsim 2$ (corresponding to a two-fold difference in strain's growth rate) or the diffusion coefficient $d \gtrsim 3$ (corresponding to a three-fold difference in strain's diffusion coefficients) coexistence was not detected for any founder density. However, the model predicted coexistence is robust over a range of differences in growth dynamics within these parameter extremes (Figure S9A and B). With strains balanced in their growth parameters, then even in the case where the weaker strain was unable to kill its competitor (c=0), coexistence remained possible at low founder densities (Figure S9C). For cases in which one strain was assumed to grow and diffuse 1.5 times as fast ($d = r = 1.5$) as its competitor, which in turn was assumed to be superior in its killing efficiency ($c > 1$), coexistence was generally observed (Figure S9D). Coexistence even occurred for extreme differences in killing coefficients ($c = 100$) at lower founder densities (Figure S9D). Finally, for low founder density, model simulations confirmed that the access to free space score was robust to changes in growth and diffusion parameters and remained an accurate predictor of competitive outcome (Figure S9E and F). These analyses show that the model hypotheses are robust to differences in strain growth dynamics.

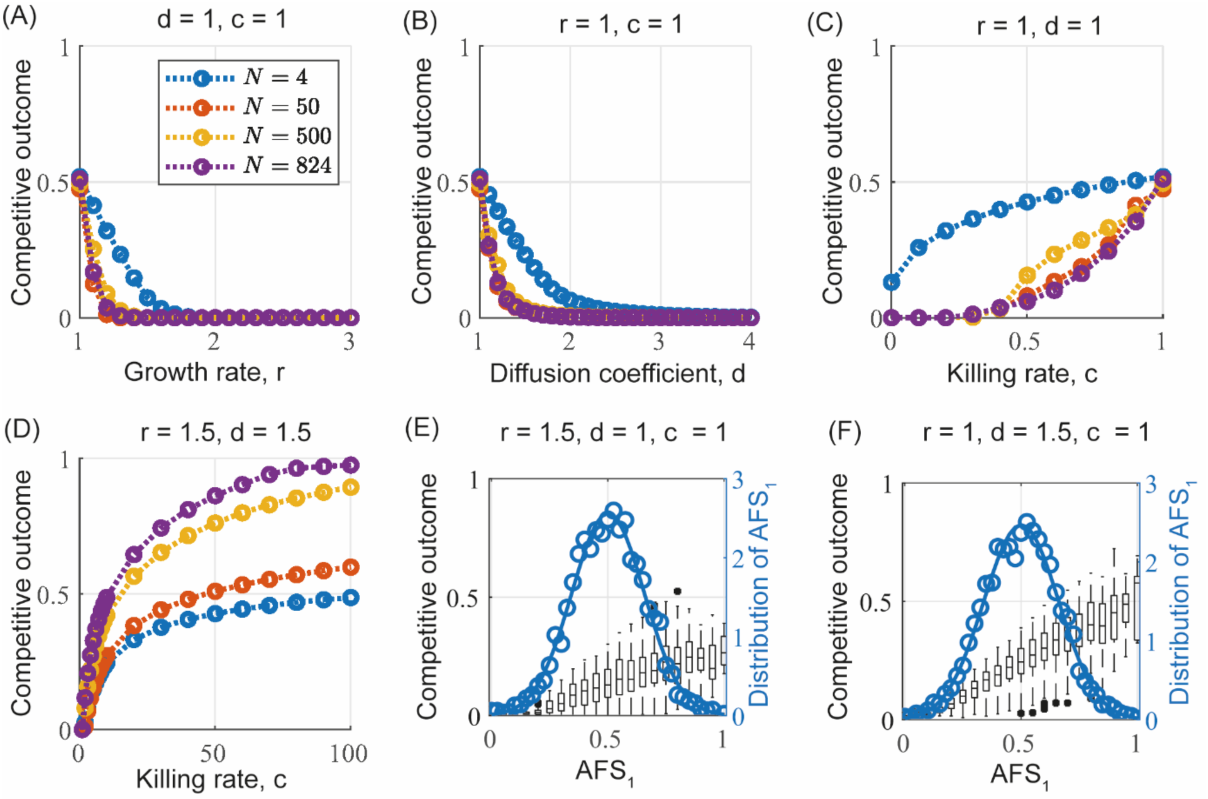

*Figure S9:* ***Impact of changes to model parameters.*** *(A-D) The relation between competitive outcome (with respect to $B_1$) and the model parameters are shown. The values of the non-varying parameters are shown. For a given founder density, all simulations are performed using identical initial conditions. (E,F) The relation between access to free space and competitive outcome is shown for two parameter sets . With the founder density fixed ($N = 6$), data are obtained from 5000 model realisations in each case. The observed probability density function for AFS is shown (circular markers) along with the density functions of fitted normal distributions ($\mu \approx 0.5, \sigma \approx 0.16$ in both) (solid lines).*

### S7 Experimental methods

#### S7.1 Growth conditions

Bacterial strains were routinely grown on lysogeny broth (LB: 1% (w/v) Bacto-peptone, 1% (w/v) NaCl, 0.5% (w/v) yeast extract and 1.5% (w/v) agar) plates either at 37°C for 18 hours or at 22°C for 64 hours or in liquid LB broth at 37°C with agitation. For biofilm formation assays, colony forming unit counts and halo formation assays the strains were grown on MSgg agar plates (5 mM potassium phosphate (pH 7), 100 mM MOPS (pH=7), 2 mM $MgCl_2$, 700 µM $CaCl_2$, 50 µM $MnCl_2$, 50 µM $FeCl_3$, 1 µM $ZnCl_2$, 2 µM thiamine 0.5% (v/v) glycerol, 0.5% (w/v) glutamate, 1.5% (w/v) agar) (2). For growth curves, MSgg media was used in a liquid form, without the addition of agar. For transforming soil isolates, a modified version of the 10 x Modified Competency (MC) media was used (10.7 g $K_2HPO_4$, 5.2 g $KH_2PO_4$, 20 g dextrose, 0.88 g sodium citrate dehydrate, 2.2 g L-glutamic acid monopotassium salt, and 1 g tryptone per 100 ml) (3). When necessary, antibiotics were used at the following concentrations: 100 µg/ml spectinomycin, 5 µg/ml chloramphenicol and 100 µg/ml ampicillin.

#### S7.2 Biofilm formation assays

Strains were streaked for single colonies on LB agar plates that were incubated at 37°C overnight. A single colony was selected to inoculate 5 ml of LB broth and grown overnight at 37°C with agitation. 100 µl of the overnight culture was placed in 3 ml of LB and grown at 37°C with agitation. Once all cultures had reached or exceeded an $OD_{600}$ of 1, incubation was stopped, and each cell culture was normalised to an $OD_{600}$ of 1.

For determining the nature of the interaction between 3610 and 6153, three independent biological replicates were performed using strains 3610 *gfp* (NRS6942), 3610 *mTagBFP* (NRS6932), 6153 *gfp* (NRS6222) and 6153 *mTagBFP* (NRS6938). Once normalised, the cultures were mixed at a 1:1 ratio in all pairwise combinations, allowing the examination of both isogenic control pairs and non-isogenic biofilm outcomes. Three technical replicated were performed for each combination and each biological repeat.

For examining the impact of different founder densities on competitive outcomes, three independent biological repeats were performed, all including isogenic mixed biofilms of both backgrounds (3610 *mTagBFP* mixed with 3610 *gfp* and 6153 *mTagBFP* mixed with 6153 *gfp*). For the non-isogenic mixes, two rounds were performed in which 3610 *gfp* was mixed with 6153 *mTagBFP* and one round was performed with the fluorescent markers swapped around (3610 *mTagBFP* mixed with 6153 *gfp*). The mixed cultures were 10-fold serial diluted and twelve technical replicates were performed for each founder density. To establish colony biofilms, 5 µl of cell culture was placed onto MSgg agar plates. Once dried, the plates were incubated at 30°C and imaged using a Leica MZ16 FA stereoscope and LAS version 2.7.1 and/ or fixed for flow cytometry after 24, 48, or 72 hours of incubation (see below). Imaging data was imported to OMERO for archiving and figure construction (4).

#### S7.3 Determining cell counts

To calculate the number of colony forming units in the culture at an $OD_{600}$ of 1, strains were grown and the $OD_{600}$ normalised as described above. The prepared cultures were serially diluted, 100 µl of the $10^{-5}$ and $10^{-6}$ dilutions spread onto MSgg agar plates and incubated at 30°C for 48 hours. Samples containing between 30 and 300 colonies were used for determining the number of colony forming units per ml of culture.

#### S7.4 Single cell analysis by flow cytometry

Biofilms grown as described above were removed from the agar plates using sterile loops, placed into 500 µl of GTA buffer (10 mM EDTA (pH=8), 20mM Tris-HCl (pH=8), 50mM glucose) and disrupted by passing through a 23-gauge needle to disrupt the biofilms before centrifuging at 17,000 x *g* for 5 minutes. The supernatant was removed, the pelleted cells were suspended in 500 µl of 4% (w/v) formaldehyde and incubated at room temperature for 7 minutes. The tubes were centrifuged at 17,000 x *g* for 5 minutes, the formaldehyde was discarded, and the samples washed by suspending the pellet in 500 µl of GTA buffer, centrifuging for 5 minutes and removing the supernatant. The pellets were resuspended in 500 µl of GTA buffer and the samples stored at 4°C. Prior to flow cytometry, the samples were mildly sonicated and 0.1 – 1 µl of the cell sample added to 1 ml of PBS buffer supplemented with 0.5% (w/v) BSA. Flow cytometry was performed using a LSRFortessa™ (BD biosciences) instrument. The results were analysed using Flowjo (version 10.7.1) to extract the % of cells expressing GFP and mTagBFP in each sample. Figures were constructed using GraphPad prism 7.

#### S7.5 Antibiosis halo assays

Strains were grown on LB agar plates. Single colonies were used to establish cultures in 5 ml of LB that were grown at 37°C with agitation overnight. 100 µl of the overnight cultures was placed in 3 ml of LB and grown at 37°C with agitation for 3 hours. At this point the cultures were normalised to a standard $OD_{600}$ of 0.5. For the lawn (target), 1 µl of each normalised culture was mixed with 100 µl of LB that spread across the surface of an MSgg plate. For the spot (attacker), 6 µl of the normalised cultures was placed in the centre of the agar where the target cells had been spread. The plates were left at room temperature for an hour to dry and incubated at 30°C. The cells were imaged after 24 and 48 hours using a using a Leica MZ16 FA stereoscope and LAS version 2.7.1. The imaging data was uploaded to OMERO (4) for figure construction.

#### S7.6 Growth curves

NCIB 3610 and NRS6153 were grown on LB agar plates at 37°C overnight to obtained single colonies. The following day, 5 ml cultures of LB broth were inoculated with single colonies and incubated at 37°C overnight with gentle agitation. 100 µl of the overnight cultures was added to 3 ml of LB broth and grown at 37 °C with agitation for 4 hours. The bacteria were harvested by centrifugation and suspended in 1 ml of MSgg media. The $OD_{600}$ was measured and normalising to an $OD_{600}$ of 1. The cell suspension was used to inoculate the growth curve cultures with a starting $OD_{600}$ of 0.01 in 25 ml of MSgg. These cultures were growth for 29 hours in a 30°C water bath and $OD_{600}$ measurements were taken every 2 hours over that period. The generation times were calculated by plotting the exponential phase of each growth curve and fitting an exponential trend-line which was used to resolve the generation time. 3 biological repeats were conducted and the average generation time for each strain was calculated. Growth curve data was plotted in GraphPad Prism 7.

### S8 *Bacillus subtilis* strains

#### S8.1 Strain construction

All *B. subtilis* strains used in this study are listed in Table S1. Plasmid pNW2303 is derived from pUC57 and carries the *mTagBFP* coding sequence (Table S2). The *mTagBFP* sequence was introduced by restriction enzyme digest to the pDR111 vector generating plasmid pNW2304 and allowing integration into the *amyE* locus. pDR111 carries the P*hyperspank* ($P_{IPTG}$) promoter and removal of the *lacI* gene by insertion of the *mTagBFP* sequence using HindIII and BamHI allows for constitutive expression. *E. coli* strain MC1061 was used for cloning purposes (*F'lacIQ lacZM15 Tn10 (tet)*).

For construction of NCIB 3610 derivatives expressing *gfpmut2* and *mTagBFP* respectively, plasmids pBL165 (containing the *gfpmut2* sequence flanked by the *amyE* locus, linked to a chloramphenicol resistance cassette) (5) and pNW2304 were integrated into the *B. subtilis* 168 genome (6). Homologous recombination at the *amyE* locus was confirmed using a potato starch assay (7). Each allele was introduced into the NCIB 3610 genome using SPP1 phage transduction (8).

For transformation of the soil isolate NRS6153 natural genetic competency was induced as described previously (3). Briefly, a day culture of each isolate was established in 2 ml of 1x MC media supplemented with 3 mM $MgSO_4$ and 875 µM $FeCl_3$. The cultures were grown for 4.5 hours at 37°C with gentle agitation. Approximately 25 µg of pBL165 or pNW2304 was added to 400 µl of cells and the reaction was incubated at 37°C with agitation for a further 90 minutes. The cells were plated onto LB agar containing the appropriate antibiotics for selection (chloramphenicol for pBL165 or spectinomycin for pNW2304). The resulting colonies were screened as described above to ensure integration at the *amyE* locus.

#### S8.2 Strain table

| **Strain** | **Relevant genotype** [a] | **Source/ construction** [b,c] |
|---|---|---|
| NCIB 3610 | Wild type, prototroph | BGSC |
| 168 | *trpC2* | BGSC |
| NRS6153 | Wild type, prototroph | (9) |
| NRS6222 | NRS6153 *amyE::*P*hyperspank- gfpmut2 (cml)* | pBL165 → NRS6153 |
| NRS6900 | 168 *trpC2 amyE::*P*hyperspank- gfpmut2 (cml)* | pBL165 → 168 |
| NRS6931 | 168 *trpC2 amyE::*P*hyperspank- mTagBFP (spec)* | pNW2304 → 168 |
| NRS6932 | NCIB 3610 *amyE::*P*hyperspank- mTagBFP (spec)* | SPP1 NRS6931 → NCIB3610 |
| NRS6938 | NRS6153 *amyE::*P*hyperspank- mTagBFP (spec)* | pNW2304 → NRS6153 |
| NRS6942 | NCIB 3610 *amyE::*P*hyperspank- gfpmut2 (cml)* | SPP1 NRS6900 → NCIB 3610 |

**Table S1: List of *B. subtilis* strains used in this study**

[a.] Antibiotic resistance cassettes are as follows; "spec" indicates a spectinomycin resistance cassette and "cml" indicates a chloramphenicol resistance cassette [b.] BSGC is the *Bacillus* genetic stock centre

[c.] The arrows show the direction of the strain construction where SPP1 phage or plasmid DNA (left of the arrow) are inserted into a recipient strain (right of the arrow)

### S9 Sequence of *mTagBFP* coding region

| ***mTagBFP* construct sequence (5'-3')** [a] |
|---|
| **AAGCTT**AAGGAGGTGATCATTAAAA*ATGAGCGAACTGATCAAAGAAAACATGCATATGAAACTGTACATG*<br>*GAAGGCACAGTCGATAACCATCACTTTAAATGCACATCAGAAGGCGAAGGCAAACCGTATGAAGGCACAC*<br>*AAACAATGAGAATCAAAGTTGTTGAAGGCGGACCGCTGCCGTTTGCATTTGATATTCTGGCAACATCATTTC*<br>*TGTATGGCAGCAAAACGTTTATCAATCATACACAAGGCATCCCGGATTTTTTTAAACAATCATTTCCGGAAG*<br>*GCTTTACATGGGAACGCGTTACAACATATGAAGATGGCGGAGTTCTGACAGCAACACAAGATACATCATTG*<br>*CAAGATGGCTGCCTGATCTATAATGTCAAAATTAGAGGCGTCAACTTTACAAGCAATGGCCCTGTTATGCAG*<br>*AAAAAAACACTGGGCTGGGAAGCATTTACAGAAACACTGTATCCGGCTGATGGCGGACTGGAAGGCAGAA*<br>*ACGATATGGCACTGAAACTGGTTGGCGGATCACATCTGATTGCAAACATCAAAACAACGTACCGCTCAAAA*<br>*AAACCGGCAAAAAATCTGAAAATGCCTGGCGTCTATTATGTCGATTATAGACTGGAACGCATCAAAGAAGC*<br>*GAACAACGAAACATATGTCGAACAACATGAAGTTGCAGTTGCGAGATATTGCGATCTGCCGTCAAAACTGG*<br>*GCCATAAACTGAATTA***GGATCC** |

**Table S2: Sequence of *mTagBFP* coding region**

[a] The restriction sites are shown in bold, the ribosome binding site and linker sequence are underlined, and the mTagBFP coding region is in italics.

### S10 Image analysis

We determine relative strain densities in competitive biofilm assays using data obtained through image analysis on the total intensity of a fluorescent signal in pixels where the signal is above the background threshold. To achieve this a Fiji/ImageJ (10, 11) macro was written (12). Since images were saved as multi-series Leica LIF files this macro relies on Bio-Formats Macro Extensions (13) to import the data. The macro can perform batch analysis of all images in a file, writing a summary table of results in CSV format as well as snapshot TIFF images showing detected biofilm regions as overlay outlines. The macro uses built-in functionality of ImageJ to detect biofilm regions, specifically: auto-thresholding using the "Triangle" method after optional background subtraction using the rolling ball / sliding paraboloid algorithm. A single large colony biofilm in the centre of the image or several smaller "sub-colonies" can optionally be detected (the former using the "Wand" tool, the latter using "Analyze Particles" with a specified size range). For the biofilm region in each image the following measurements are made for 2 channels: area, basic intensity statistics (mean, maximum, standard deviation, total intensity) and some colocalization statistics (Pearsons Correlation Coefficient, thresholded "Object Pearsons" (14) and optionally Manders M1 and M2 coefficients (15). Finally, percentage area within the biofilm region that is above background for each channel is reported, as well as total "foreground" signal (i.e. total signal in pixels that have above-background intensity values). These final two measurements rely on a background intensity parameter for each channel that distinguishes positive expression of the label from background.
For each image, the relative strain density was calculated by dividing the strain's foreground signal by the total foreground signal in the image.

#### S10.1 Access to free space

To calculate the access to free space score from experimental data, we used the same method as in our classification of the model initial conditions. We chose the reference circle to be of radius $R \approx 12\ mm$. The radius is an approximation as we used the number of pixels as a proxy unit for length and pixel sizes slightly varied between images. The access to free space score in our experimental assays was calculated based on images taken after 24h of incubation. Only cultures in which microcolonies did not overlap at this time point are analysed. For each image, two subimages showing the signal of one fluorescent signal only were converted into binary images using colour segmentation based on Delta E colour difference. For each pixel of the image, these binary images provided information on which strain (if any) occupies the space shown in the pixel. This analysis determines the locations of the microcolonies and the access to free space score was calculated by the same means as for the mathematical model ((16), see also Section S4.2).
